# Extracellular adenosine enhances the ability of PMNs to kill *Streptococcus pneumoniae* by inhibiting IL-10 production

**DOI:** 10.1101/716456

**Authors:** Nalat Siwapornchai, James N. Lee, Essi Y. I. Tchalla, Manmeet Bhalla, Jun Hui Yeoh, Sara E. Roggensack, John M. Leong, Elsa N. Bou Ghanem

**Author notes:** These authors contributed equally to this work (Nalat Siwapornchai, James N. Lee). Summary Sentence: Extracellular adenosine produced by CD73 promotes the ability of PMNs to kill *Streptococcus pneumoniae* by blunting IL-10 production. Corresponding author information: Elsa N. Bou Ghanem, 955 Main Street, Buffalo, NY, 14203.

## Abstract

PMNs are crucial for initial control of *Streptococcus pneumoniae* (pneumococcus) lung infection; however, as the infection progresses their persistence in the lungs becomes detrimental. Here we explored why the anti-microbial efficacy of PMNs declines over the course of infection. We found that the progressive inability of PMNs to control infection correlated with phenotypic differences characterized by a decrease in CD73 expression, an enzyme required for production of extracellular adenosine (EAD). EAD production by CD73 was crucial for the ability of both murine and human PMNs to kill *S. pneumoniae.* In exploring the mechanisms by which CD73 controlled PMN function, we found that CD73 mediated its anti-microbial activity by inhibiting IL-10 production. PMNs from wild type mice did not increase IL-10 production in response to *S. pneumoniae,* however, CD73^-/-^ PMNs up-regulated IL-10 production upon pneumococcal infection *in vitro* and during lung challenge. IL-10 inhibited the ability of wild type PMNs to kill pneumococci. Conversely, blocking IL-10 boosted the bactericidal activity of CD73^-/-^ PMNs as well as host resistance of CD73^-/-^ mice to pneumococcal pneumonia. CD73/IL-10 did not affect apoptosis, bacterial uptake and intracellular killing or production of anti-microbial Neutrophil Elastase and Myeloperoxidase. Rather, inhibition of IL-10 production by CD73 was important for optimal ROS production by PMNs. ROS contributed to PMN anti-microbial function as their removal or detoxification impaired the ability of PMNs to efficiently kill *S. pneumoniae*. This study demonstrates that CD73 controls PMN anti-microbial phenotype during *S. pneumoniae* infection.

## 1. Introduction

Neutrophils, also known as polymorphonuclear leukocytes or PMNs, play a major role in host defense against *S. pneumoniae* lung infection (1, 2). PMNs are required to control bacterial burden early in the infectious process. In mouse models, early PMN recruitment into the lungs within the first 12 hours following pneumococcal pulmonary infection coincided with a decrease in bacterial numbers and immunodepletion of PMNs prior to infection resulted in host lethality (3). *Ex vivo*, PMNs are thought to kill *S. pneumoniae* by engulfing the bacteria and producing serine proteases including cathepsin G (CG) and neutrophil elastase (NE) (4), which also play a role in controlling bacterial numbers in murine models of pneumonia (5). Interestingly, later during infection, PMN persistence in the lungs promotes disease (3, 6, 7). In fact, we previously found that depletion of PMNs 18 hours after pneumococcal lung infection results in reduced bacterial numbers and enhanced mouse survival (3). These findings demonstrate that while PMNs are required at the start of infection, their persistence is detrimental for host survival. This also suggested that the antibacterial function of these innate immune cells and their ability to clear pneumococci is altered during the course of infection (3). However, the mechanisms of this alteration in PMN phenotype remain unexplored.

A major regulator of host resistance to pneumococcal infection is extracellular adenosine (EAD) (3). Upon damage due to a variety of insults including infection, ATP is thought to leak from damaged cells into the extracellular space and is converted into EAD by the sequential action of two exonucleosidases, CD39 and CD73 (8). EAD can then signal via four known receptors, i.e. A1, A2A, A2B and A3 (9). Several drugs targeting this pathway are in clinical studies (9), making it an attractive avenue for modulating immune responses during infection. We previously found that blocking EAD production by CD73 dramatically increases bacterial numbers in organs and results in host lethality upon *S. pneumoniae* lung infection in mice (3). EAD production by CD73 regulates PMN recruitment to the lungs and potentially boosts PMN function (3).

IL-10 is an anti-inflammatory cytokine associated with impaired control of pneumococcal pneumonia (10, 11). In mouse models, IL-10 levels increase in the lungs within 12 hours following pneumococcal challenge (10). Intranasal administration of IL-10 at the time of bacterial challenge results in decreased host survival and increased bacterial numbers in the lungs and blood, while blocking IL-10 enhances bacterial clearance and boosts host survival (10). Interestingly, while IL-10^-/-^ mice have reduced lung and systemic bacterial loads, these mice suffer increased mortality due to excessive pulmonary inflammation (12). IL-10 can be produced by many types of immune cells and it is now appreciated that murine PMNs produce this anti-inflammatory cytokine during infections (13–16). PMNs are a significant source of IL-10 in sepsis models (17) and stimulation with several bacterial pathogens, including *Escherichia coli*, *Shigella flexneri* and mycobacterial BCG, triggers IL-10 production by PMNs both *in vitro* and *in vivo* within 8 hours of infection (16). Parasitic infections such as *Leishmania major* (14) and *Trypanosoma cruzi* (15) also induce IL-10-producing PMNs during mouse infection. IL-10 production by PMNs may reduce their microbicidal capacity, as *in vitro* treatment of PMNs with IL-10 impairs their ability to phagocytose *Staphylococcus aureus*, *Candida albicans* and *E. coli* and blunts superoxide production (18, 19). Further, in a *S. aureus* burn infection model, a subset of IL-10 producing PMNs is associated with impaired host resistance to infection (13). It is not currently known if *S. pneumoniae* infection triggers IL-10 production by PMNs.

In this study, we explored the involvement of CD73 in the diminished anti-microbial efficacy of PMNs during the course of pneumococcal pneumonia. We found that during the course of lung infection, CD73 expression on PMNs decreased. Notably, this change in CD73 expression coincided with an increase in IL-10-producing PMNs. CD73 suppressed IL-10 production by PMNs during infection and was required for the ability of these cells to kill *S. pneumoniae.* This study identifies a role for CD73 in maintaining the antimicrobial phenotype of PMNs and elucidates the mechanisms by which CD73 regulates PMN anti-microbial function and host resistance against *S. pneumoniae*.

## 2. Materials and Methods

### 2.1 Mice

All experiments were conducted in accordance with Institutional Animal Care and Use Committee (IACUC) guidelines. Wild type (WT) C57BL/6 mice were purchased from Jackson Laboratories (Bar Harbor, ME). CD73^-/-^ mice on C57BL/6 background (8) were purchased from Jackson Laboratories and bred at a specific-pathogen free facility at Tufts University and The University at Buffalo. Female 8-12-week-old mice were used in all experiments.

### 2.2 Bacteria

Wild type *S. pneumoniae* TIGR4 AC316 strain (serotype 4) and pneumolysin-deletion mutant (Δ*PLY*) *S. pneumoniae* (20) were kind gifts from Andrew Camilli. The superoxide dismutase in-frame deletion mutant (Δ*sodA*) strain was constructed via allelic exchange using linear pieces of DNA amplified by PCR and the co-transformation approach previously outlined (21). Primers SP_0766_F1 and SP_0766_R1 (table 1) were used to generate a linear piece of DNA with homologous sequences in the pneumococcal genome upstream of *sodA*, and primers SP_0766_F2 and SP_0766_R2 (table 1) amplified the region downstream of *sodA*. These primers were designed with overlapping sequences, such that when assembled using NEBuilder followed by a final PCR amplification with the F1 and R2 primers (table 1), the entirety of the *sodA* sequence is deleted except for the start and stop codons. The ΔSP_1051::cat sequence was amplified from an existing ΔSP_1051::cat strain in the AC316 background (21) using primers OS38 and OS39 (table 1). The two pieces of linear DNA were transformed simultaneously into AC316 in a molar ratio of unmarked to marked approximately 20:1. In cells that took up both pieces of linear DNA and recombined them into the genome, *sodA* was deleted from the genome while neutral gene SP_1051 was replaced with a chloramphenicol resistance cassette. Chloramphenicol-resistant transformants were screened for the gene deletion via colony PCR using primers SP_0766_ColonyPCRF and SP_0766_ColonyPCRR (table 1). Double-positive clones were grown and saved in THY/glycerol for further verification. PCR amplification of both the ΔSP_1051::cat and Δ*sodA* regions were performed using template gDNA extracted from a double-positive clone. Amplified DNA was run on agarose gels and imaged under UV-illumination. All bacteria were grown at 37°C in 5% CO_2_ in Todd-Hewitt broth supplemented with 0.5% yeast extract and oxyrase until culture reached mid-exponential phase. Bacterial aliquots were frozen at −80°C in the growth media with 20% (v/v) glycerol. Prior to use, aliquots were thawed on ice, washed and suspended in PBS to obtain the appropriate concentration. The inoculums were then confirmed by serial dilution and dribble plating on Tryptic Soy Agar plates supplemented with 5% sheep blood agar.

**Table I.**
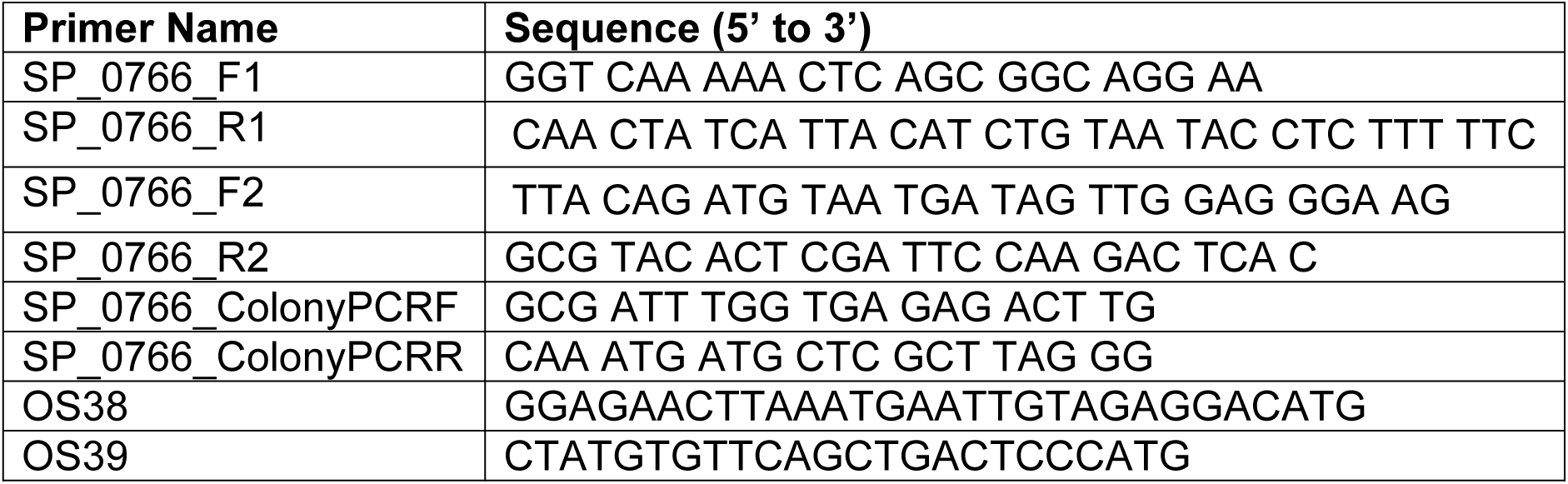
Primers used for generation of *ΔsodA S. pneumoniae*

### 2.3 Isolation of PMNs

Bone marrow cells were harvested from femurs and tibias of uninfected mice, flushed with RPMI 1640 supplemented with 10% FBS and 2 mM EDTA, and resuspended in PBS. Neutrophils were then separated from the rest of the bone marrow cells through density gradient centrifugation, using Histopaque 1119 and Histopaque 1077 as previously described (22). The isolated neutrophils were resuspended in Hanks’ Balanced Salt Solution (HBSS)/0.1% gelatin without Ca^2+^ and Mg^2+^, and used in subsequent assays. The purity of neutrophils was also measured by flow cytometry using APC-conjugated anti-Ly6G and 85-90% of enriched cells were Ly6G+.

### 2.4 Opsonophagocytic killing assay

The ability of PMNs to kill pneumococci was assessed *ex vivo* through an Opsonophagocytic (OPH) killing assay as previously described (3, 23). Briefly, 100μl reactions in HBSS/0.1% gelatin consisted of 1×10^5^ PMNs incubated with 1×10^3^ bacteria grown to mid log phase and pre-opsonized with 3% mouse sera. The reactions were incubated rotating for 45 minutes at 37°C. Where indicated, PMNs were incubated with adenosine (100μM), rIL-10 (50ng/mL), anti-IL10 (1ug/mL JES5-2A5) or isotype control (1ug/mL), 1x protease inhibitor cocktail, the CD37 inhibitor (100μM α,β methylene ADP), the CD39 inhibitor (100μM POM-1), the ROS inhibitor Diphenyleneiodonium (DPI,10μM) or the anti-oxidants EUK134 (25μM) or Ascorbic acid (100μM) for 30 minutes prior to adding pre-opsonized bacteria. IL-10 reagents were purchased from Biolegend and the inhibitors from Sigma. HBSS with 3% sera was added to PMNs without treatment as the control. Percent killing was determined by dribble plating on blood agar plates and calculated in comparison to no PMN control under the exact same conditions (+/-treatments).

### 2.5 IL-10 Enzyme-Linked Immunosorbent Assay (ELISA)

1×10^6^ Bone marrow PMNs were incubated with HBSS/0.1% gelatin or adenosine for 30 minutes, followed by 45 minutes infection at 37° C with pre-opsonized *S. pneumoniae* TIGR4 at a MOI of 2 or mock treated with HBSS and sera only (uninfected). The reactions were spun down and the supernatants were obtained to measure IL-10 production using Mouse IL-10 ELISA kit (eBioscience) according to manufacturer’s instructions.

### 2.6 ROS Assay

Following isolation from the bone marrow, PMNs were re-suspended in HBSS (Ca^2+^ and Mg^2+^ free) and acclimated at room temperature for one hour. The cells were then spun down and re-suspended in KRP buffer (Phosphate buffered saline with 5mM glucose, 1mM CaCl_2_ and 1mM MgSO_4_) and equilibrated at room temperature for 30 minutes. The cells were then seeded in 96-well white LUMITRAC^TM^ plates (Greiner Bio-One) at 5×10^5^ PMNs per well, treated with adenosine, anti-IL-10 or rIL-10 for 30 minutes at 37° C. For detection of extracellular ROS, 50μM Isoluminol (Sigma) plus 10U/ml HRP (Sigma) were added, while for detection of intracellular ROS, 50μM Luminol (Sigma) was added to the wells as previously described (24–27). The cells were infected with pre-opsonized *S. pneumoniae* TIGR4 at a MOI of 2 or mock treated with buffer containing 3% mouse sera (uninfected). PMNs treated with 100 nM Phorbol 12-myristate 13-acetate (PMA) (Sigma) were used as a positive control. Luminescence was immediately read (following infection) over a period of one hour at 37° C in a pre-warmed Biotek Plate reader. Wells containing buffer and Isoluminol plus HRP or Luminol alone were used as blanks.

### 2.7 Animal infections and scoring

Mice were lightly anesthetized with isoflurane and challenged intra-tracheally (i.t.) with 5×10^5^ colony-forming units (CFU) of WT *S. pneumoniae* TIGR4 strain by instilling 50μl of bacteria directly into the trachea with the tongue pulled out to facilitate delivery of bacteria directly into the lungs (3). Following the infection, mice were monitored daily for weight loss and blindly scored for signs of sickness including weight loss, activity level, posture and breathing, scored as healthy [0] to severely sick [21] as previously described (28). The lung and brain were harvested and homogenized in sterile PBS. Blood was collected to follow bacteremia. Each sample was diluted in sterile PBS and dribble plated on blood agar plates to enumerate bacterial numbers.

### 2.8 Isolation of cells from the lungs and flow cytometry

Mice were perfused with 10ml PBS, the lungs removed, washed in PBS, and minced into small pieces. The lungs were then digested for 1 hour with RPMI 1640 supplemented with 10% FBS, 1 mg/ml Type II collagenase (Worthington), and 50 U/ml Deoxyribonuclease I (Worthington) at 37° C/ 5% CO_2_. Single-cell suspensions were obtained by mashing the digested lungs, and the red blood cells were removed by treatment with a hypotonic lysis buffer (Lonza). Cells were analyzed using flow cytometry. Intracellular cytokine staining (ICS) was performed using the Cytofix/Cytoperm kit (BD Biosciences). GolgiPlug was added to the digestion media and following red blood cell lysis, the cells were incubated with RPMI 1640 supplemented with 10% FBS and Golgi Plug for 3 more hours at 37° C/ 5% CO_2_. Cells were surface stained with anti-mouse CD45 (clone 30-F11, eBioscience), Ly6G (clone 1A8, BD Biosciences) and CD73 (Clone TY/11.8, eBioscience). For intracellular staining, cells were permeabilized and stained with IL-10 (clone JES5-16E3, eBioscience) or isotype control (Rat IgG2b Κ Isotype Control, Biolegend). Fluorescence intensities were measured on a FACSCalibur and at least 25,000 events for lung tissue were analyzed using FlowJo.

### 2.9 Adoptive transfer of PMNs

Bone marrow PMNs were isolated from uninfected mice using density gradient centrifugation as described above and resuspended in PBS. Mice either received 2.5×10^6^ cells via intraperitoneal (i.p.) injection as previously described (29) or mock-treated (PBS). One hour post-transfer, mice were challenged i.t. with 5×10^5^ CFU of *S. pneumoniae.* Mice were euthanized at 24 hours post infection, and the lungs, brain, and blood were collected and plated on blood agar plates for CFU. The presence of transferred PMNs in the circulation of recipient mice was confirmed by flowcytometry (Fig S1).

**Figure 1.**
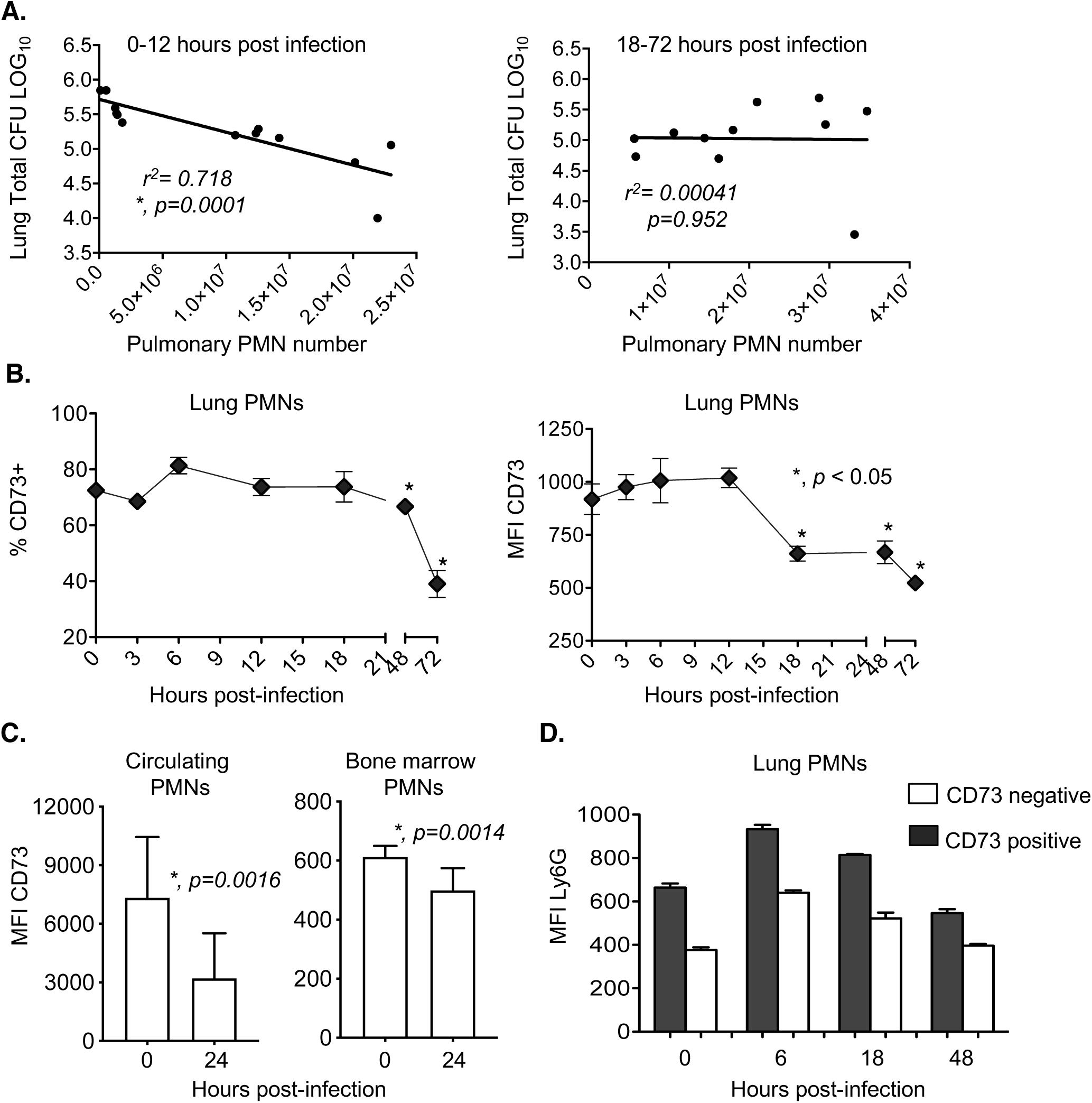
Expression of CD73 on pulmonary PMNs changes over the course of infection. C57BL/6 mice were inoculated i.t with 5×10^5^ CFU of *S. pneumoniae* TIGR4. The blood, bone marrow and lungs were harvested, plated for bacterial enumeration and analyzed by flow cytometry. (A) Mice were euthanized at 0, 6, 12, 18, 24 and 48 hours following infection. We then compared bacterial numbers in the lungs to PMN numbers (Ly6G+ cells) for the first 0-12 and subsequent 18-72 hours post infection and performed Pearson correlation analysis where asterisks denote significant correlation. We gated on PMNs (Ly6G+ cells) and also monitored (B) the percentage of cells expressing CD73 and (B-C) the amounts of CD73 (mean fluorescent intensity or MFI) in the indicated organs, at the indicated time points. (D) We gated on all Ly6G+ cells and then gated on CD73 positive versus negative populations and compared the expression (MFI) of Ly6G. Data are pooled from two separate experiments with 4-5 mice per time point (A) and 4-7 mice per time point (B-D). Asterisks indicate significant differences from uninfected controls calculated by Student’s t-test.

### 2.10 Blocking IL-10 in vivo

Mice were treated with 0.1mg/mouse of IL-10 blocking antibody (JES52A5, Biolegend) or the isotype control (Rat IgG1 Κ Isotype Control, Biolegend) through i.p. injection. After 2 hours, mice were then challenged i.t. with 5×10^5^ CFU of *S. pneumoniae.* The lungs and blood were collected two days post-infection to determine bacterial burden.

### 2.11 Assays with human PMNs

Young (23-38 years old), male and female healthy human volunteers were recruited in accordance with The University at Buffalo Human Investigation Review Board (IRB) and signed informed consent forms. Individuals taking medication, reporting symptoms of infection within the last 2 weeks or that were pregnant were excluded from the study. Whole blood was obtained using acid citrate/dextrose as an anti-coagulant. PMNs were isolated using a 2% gelatin sedimentation technique as previously described (30) which allows for isolation of active PMNs with ∼90% purity. 5 x 10^5^PMNs were treated with the selective and competitive inhibitor of CD73, α,β methylene ADP (1 or 10μM) or DMSO vehicle control for 30 minutes at 37° C. The PMNs were then infected with 10^3^ CFU *S. pneumoniae* pre-opsonized in 10% (v/v) baby rabbit serum (Pel-Freeze) for 45 minutes at 37°C with rotation. Samples were then placed on ice to stop the process of opsonophagocytosis followed by serial dilution and plating on blood agar plates to enumerate viable CFU. The percentage of bacterial killing was calculated relative to no PMN controls incubated under the same treatment conditions.

### 2.12 mRNA measurement

RNA was extracted from 2×10^6^ human PMNs using the RNeasy Mini Kit (Qiagen) as per manufacturer’s protocol. TURBO DNA-free kit (Invitrogen) was used to digest DNA from the RNA samples prior conversion into cDNA. RNA concentration and 260/280 ratio were determined using Nano-drop (Thermo Fischer Scientific). For each sample, 500ng of RNA was converted into cDNA using SuperScript VILO^TM^ cDNA synthesis kit (Life Technologies) according to the manufacturer’s protocol. RT-PCR was performed using CFX96 Touch™ Real-Time PCR Detection System from Bio-Rad and CT (cycle thresh-hold) values were determined using the following TaqMan probes from Life Technologies (Thermo Fischer Scientific): GAPDH (human Hs99999905_m1) and IL-10 (human Hs00961622_m1). Each sample was run in duplicates. Data were analyzed by the comparative threshold cycle (2^-ΔCT^) method, normalizing the CT values obtained for IL-10 expression to those for GAPDH of the same sample. For comparison of IL-10 levels upon infection, relative quantity of transcripts (RQ) values were calculated by the ΔΔCT method by using the formula RQ =2^^_^ (ΔΔCT) (31). The ΔΔCT values were obtained by subtracting ΔCT value of the infected (test) from that of the uninfected control.

### 2.13 Statistics

All statistical analysis was performed using Prism7 (Graph Pad). CFU data were log-transformed to normalize distribution. For all graphs, the mean values +/-SD are shown. Significant differences were determined by 2-way ANOVA followed by Sidak’s multiple comparisons test or Student’s t-test where *p* values less than 0.05 were considered significant (as indicated by asterisks). Pearson test was used to determine correlation.

## Online Supplemental Material

### Adenosine measurement

Blood was collected from mice by venipuncture in microtainer tubes. The tubes were centrifuged at 9000rpm for 2 minutes and the sera obtained. Adenosine level in the sera was measured using the Adenosine Assay Fluorometric Kit (BioVision) as per manufacturer’s instructions.

### Bacterial uptake and intracellular killing assay

To determine uptake/phagocytosis and intracellular killing by PMNs, we performed a gentamicin-protection assay. Isolated PMNs from uninfected mice were incubated with pre-opsonized wild type or Δ*PLY S. pneumoniae* (MOI=2) for 10 minutes at 37°C. Gentamycin (100ug/ml) was then added for 40 minutes to kill extracellular bacteria. The reactions were washed three times with HBSS and resuspended in 100ul HBSS/0.1% gelatin. To measure initial bacterial uptake, the reactions were diluted and plated on blood agar plates and the percentage of the input inoculum that was engulfed was calculated. To determine intracellular killing, the reactions were returned to 37°C and incubated for 30 minutes and 90 minutes. The reactions were diluted and plated on blood agar plates and the percentage of the engulfed inoculum (at 10 minutes) that was killed was then calculated.

### Myeloperoxidase (MPO) levels, Neutrophil Elastase and Cathepsin G activity assays

1×10^6^ Bone marrow PMNs were treated with adenosine, anti-IL-10 or rIL-10 or mock treated with HBSS/0.1% gelatin for 30 minutes at 37° C. The PMNs were then infected with pre-opsonized *S. pneumoniae* TIGR4 at a MOI of 2 or mock treated with buffer containing 3% mouse sera (uninfected) for 45 minutes at 37°C. The reactions were spun down and the supernatants were used to measure neutrophil elastase and cathepsin G activities as well as MPO levels with the Neutrophil Elastase Activity Assay Kit Fluorometric (Abcam), the Cathepsin G Activity Assay Kit (Abcam) and MPO ELISA (Invitrogen) respectively (30). The cells were lysed with lysis buffer containing 1% Triton-X, 50mM Tris, 0.1% SDS, 0.5% sodium deoxycholate and 150mM NaCl. MPO levels in the pellets were also measured by ELISA.

### Apoptosis Assay

PMNs were pre-treated with buffer or incubated with anti-IL-10 or rIL-10 for 30 minutes at 37° C then infected with pre-opsonized *S. pneumoniae* TIGR4 at a MOI of 2 or mock treated with buffer containing 3% mouse sera (uninfected) for 45 minutes at 37° C. The percentage of apoptotic cells were then determined by flow cytometry using the FITC Annexin V apoptosis detection kit with PI (BioLegend) following manufacturer’s instructions.

### Tracking of transferred PMNs

Bone marrow PMNs were isolated from uninfected CD73^-/-^ or wildtype mice using density gradient centrifugation and then labeled with CellTracker Green (CMFDA (5-Chloromethylfluorescein Diacetate) and CellTracker Orange (CMTMR (5-(and-6)-4-Chloromethyl Benzoyl Amino Tetramethylrhodamine) respectively as previously described (22). CD73^-/-^ mice were then injected with a 1:1 ratio of PMNs from each strain (1×10^6^ cells) via intraperitoneal (i.p.) injection. Three hours post transfer, recipient mice were euthanized, blood collected and the presence of transferred cells assessed by flow cytometry.

## 3. Results

### 3.1 CD73 expression on PMNs decreases over the course of infection

We previously found that PMNs are required for protection at the beginning of *S. pneumoniae* lung infection (3), but are detrimental at later times. To test if there was a correlation between PMN numbers in the lungs and bacterial burden, we infected young C57BL/6 mice intra-tracheally (i.t.) with 5×10^5^ colony-forming units (CFU) of *S. pneumoniae* TIGR4 strain. We then monitored the bacterial burden and pulmonary influx of PMNs and performed correlation analysis between PMN number and lung CFU for 0-12 hours vs. 18-72 hours following infection. We found that PMN influx into the lungs strongly correlated with a decrease in bacterial burden for the first 12 hours following infection (Fig 1A left panel, R-squared =0.71, *p=0.0001*). However, there was no correlation between PMN presence in the lungs and bacterial numbers for the remainder of the infection (Fig 1A right panel) suggesting that PMNs are no longer able to control the infection.

To determine if this was associated with a difference in the phenotype of pulmonary PMNs over time, we monitored the expression of the EAD-producing enzyme CD73 on PMNs in the lungs following challenge. We found that in the first 18 hours of infection, the majority (∼75-80%) of PMNs in the lungs expressed CD73 (Fig. 1B). However, both the percentage of PMNs expressing CD73 and the amount (measured by the mean fluorescent intensity or MFI) of CD73 expressed on PMNs in the lungs significantly dropped over the course of infection. CD73 expression had dropped to half by 72 hours and the drop started to occur after 12 hours post-infection, a time point after which PMN presence in the lungs no longer correlated with control of bacterial numbers (Fig 1A) (3). The decrease of CD73 expression also occurred on circulating PMNs and bone marrow PMNs (Fig. 1C), suggesting that this was a systemic decrease and not only localized in the lungs. A recent study identified a population of PMNs in the spleen with intermediate expression of Ly6G on their surface that exhibited a lowered ability to engulf *S. pneumoniae* during i.v. infection (32). When we compared Ly6G expression, we found that CD73 positive pulmonary PMNs had significantly higher expression of Ly6G (MFI) on their surface at all the time points tested (Fig. 1D). These data show that there are changes in PMNs over the course of pneumococcal pneumonia and that low CD73 expression on PMNs may be indicative of a PMN subset that is associated with lower anti-microbial activity.

### 3.2 CD73 and extracellular adenosine are required for the ability of PMNs to kill *S. pneumoniae*

We previously found that EAD production and signaling is required for protection against *S. pneumoniae* lung infection (3, 33). To test if EAD production has a role in anti-pneumococcal function of PMNs, we compared *ex vivo* opsonophagocytic killing of pneumococci by PMNs isolated from the bone marrow of CD73^-/-^ or wild type C57BL/6 (WT) mice. Strikingly, CD73^-/-^ PMNs completely failed to kill *S. pneumoniae.* In fact, the presence of these PMNs promoted an increase in bacterial numbers as indicated by negative bacterial killing on the graph (Fig. 2A). To test if the inability of CD73^-/-^ PMNs to kill bacteria was due to a defect in EAD production, we added adenosine to the opsonophagocytic reactions. Adenosine levels in the sera of WT mice varied, but on average increased five-fold to 52.6 +/-29.0 μM upon pneumococcal infection as measured by a fluorometric kit (Fig S 2A). We found that supplementing CD73^-/-^ PMNs with EAD at 100 μM (closer to the higher levels found in infected mice) restored the ability of these cells to kill bacteria back to levels comparable with WT PMNs. Addition of exogenous EAD had no significant effect on bacterial killing by WT PMNs (Fig 2A). However, when we pharmacologically inhibited CD39 or CD73, the enzymes required for sequential dephosphorylation of extracellular ATP to EAD, we observed a significant reduction in the ability of WT PMNs to kill *S. pneumoniae* (Fig 2B). We confirmed that the difference in killing observed was not due to difference in bacterial survival in WT versus CD73^-/-^ sera or direct toxic effects of EAD (Fig. S2B and C) or the inhibitors on bacterial viability as previously described (3).

**Figure 2.**
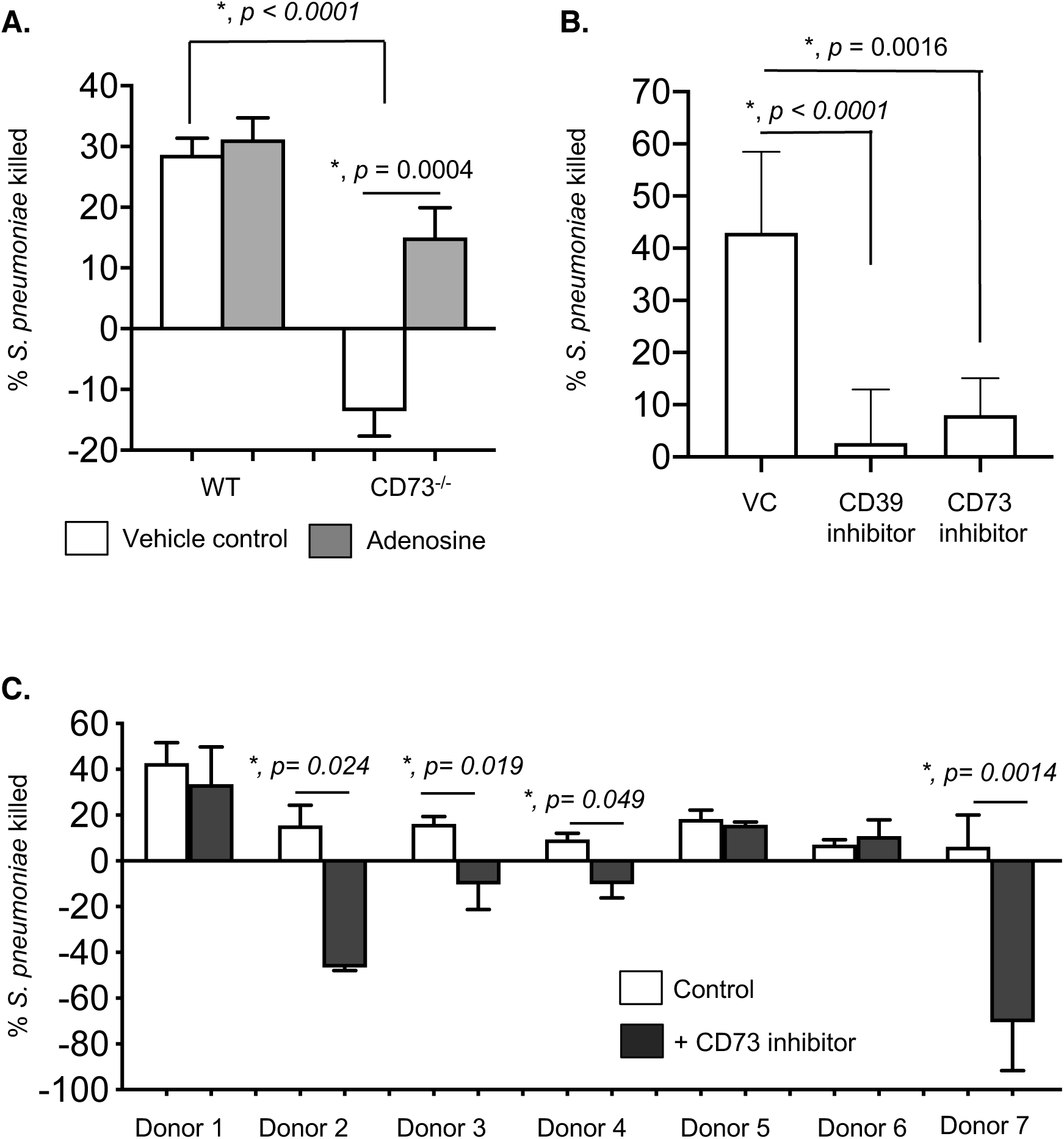
CD73 and extracellular adenosine are required for the ability of neutrophils to kill *S. pneumoniae.* (A) PMNs were isolated from the bone marrow of C57BL/6 (WT) or CD73^-/-^ mice and treated with 100μM Adenosine or PBS (vehicle control) for 30 minutes at 37°C. (B) PMNs were isolated from the bone marrow of C57BL/6 (WT) mice and treated with 100μM of the CD37 inhibitor (α,β methylene ADP), 100μM of the CD39 inhibitor (POM-1) or PBS (vehicle control-VC) for 30 minutes at 37°C. The reactions were then infected with *S. pneumoniae* pre-opsonized with homologous sera for 45 minutes at 37°C. Reactions were then stopped by placing samples on ice and viable CFU were determined after serial dilution and plating. The percentage of bacteria killed upon incubation with PMNs was determined by comparing surviving CFU to a no PMN control. Positive percent killing indicates bacterial death while negative percent indicates bacterial growth. Data shown are pooled from three separate experiments (n=3 biological replicates or mice per strain) where each condition was tested in triplicate (n=3 technical replicates) per experiment. Asterisks indicate significance calculated by Student’s t-test. (C) PMNs were isolated from the blood of young healthy donors and pre-treated with the CD73 inhibitor (α,β methylene ADP) or vehicle control for 30 minutes at 37°C and then incubated for 45 minutes with complement-opsonized *S. pneumoniae* TIGR4. For each donor, the average percent bacterial killing compared to a no PMN control was calculated from triplicate wells per condition. Data from 7 donors are shown. Significant differences denoted by asterisk, were determined by paired t-test.

To test the clinical relevance of our findings, we tested the effect of CD73 inhibition on the anti-bacterial activity of PMNs from human donors. PMNs were isolated from the blood of young healthy donors and treated with the CD73 inhibitor or vehicle control *in vitro* and their ability to kill pneumococci was assessed. We found that in over half (i.e. 4 out of 7) of the donors tested, CD73 inhibition completely abrogated the ability of PMNs to kill pneumococci and in fact promoted bacterial growth in the presence of PMNs instead (Fig. 2C). These findings demonstrate that production of EAD by CD73 is required for the ability of PMNs to kill pneumococci.

### 3.3 Adoptive transfer of PMNs from wild type mice boosts resistance of CD73^-/-^ mice to S. *pneumoniae*

To test the role of EAD production by PMNs *in vivo*, we adoptively transferred 2.5×10^6^ PMNs isolated from WT or CD73^-/-^ mice into CD73^-/-^ mice, one hour later infected the mice with 5×10^5^ CFU of *S. pneumoniae* i.t. and then compared bacterial burdens in the lung, blood and brain at 24 hours post infection. We found that transfer of WT PMNs significantly reduced the systemic spread of pneumococci and resulted in a 50 and 5-fold reduction in blood and brain bacterial loads respectively (Fig. 3B and C) when compared to no transfer controls. In fact, bacteremia in CD73^-/-^ mice that had received WT PMNs was comparable to that observed in WT controls (Fig 3B). Transfer of WT PMNs had no significant effect on bacterial numbers in the lungs of CD73^-/-^ recipients (Fig. 3A). Surprisingly, we found that adoptive transfer of CD73^-/-^ PMNs worsened susceptibility to infection and resulted in a ∼10-fold increase in bacterial numbers in all sites (lung, blood, brain) tested (Fig. 3A-C) in CD73^-/-^ recipients when compared to no transfer controls. WT mice that received CD73^-/-^PMNs also had a slight (but not statistically significant) increase in pulmonary bacterial numbers (Fig. 3A). This was consistent with our *in vitro* findings where bacterial numbers increased in the presence of CD73^-/-^ PMNs. These findings suggest that CD73 expression by PMNs is required and sufficient to promote resistance to *S. pneumoniae* infection.

**Figure 3.**
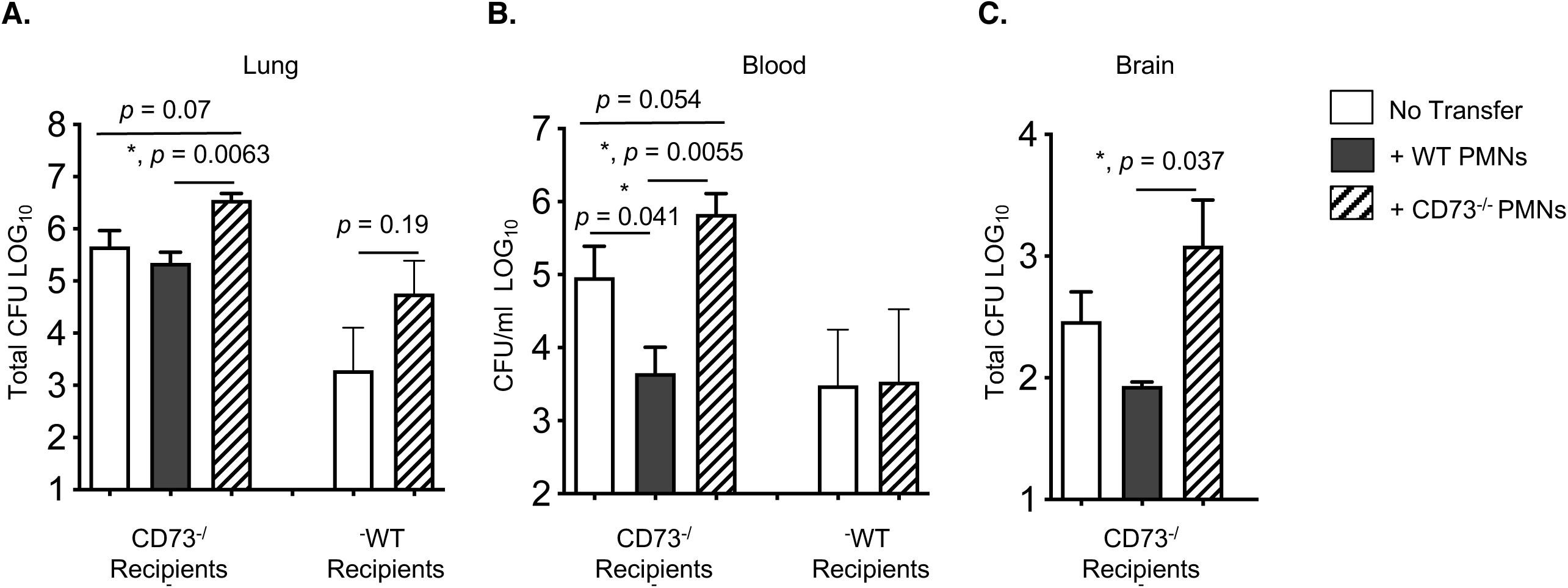
Adoptive transfer of PMNs from wild type mice boosts resistance of CD73^-/-^ mice to *S. pneumoniae.* C57BL/6 (WT) or CD73^-/-^ mice were mock treated (no transfer) or injected i.p with 2.5×10^6^ of the indicated PMNs isolated from the bone marrow of C57BL/6 or CD73^-/-^ mice. One hour post transfer, mice were infected i.t with 5×10^5^ CFU of *S. pneumoniae* and bacterial numbers in the lung (A), blood (B) and brain (C) were determined 24 hours post infection. Significant differences, determined by Student’s t-test, are indicated by asterisks. Pooled data from n=5 mice per group are shown.

### 3.4 Extracellular adenosine blocks IL-10 production by PMNs following *S. pneumoniae* infection

EAD was previously shown to play a role in regulating production of cytokines by immune cells (34) including IL-10 by macrophages (35–37). IL-10 is an anti-inflammatory cytokine whose production early on during pneumococcal infection was reported to be detrimental for host resistance (10). To test whether EAD was targeting IL-10, we compared IL-10 production by WT and CD73^-/-^ PMNs upon *in vitro* infection with *S. pneumoniae.* We found that at baseline, PMNs from both mice strains produced some IL-10 but there was no significant difference in levels. However, upon infection, within the 45 minute timeframe of our *in vitro* assays, CD73^-/-^ PMNs produced significantly more IL-10 as compared to WT PMNs (Fig. 4A). Further, while WT PMNs did not upregulate IL-10 production upon pneumococcal infection, CD73^-/-^ PMNs displayed a ∼ 2.5-fold increase in IL-10 above resting baseline levels (Fig 4A and B). Importantly, this upregulation was significantly blunted upon exposure to extracellular adenosine (Fig. 4B). When we examined PMNs isolated from three human donors, we could not detect any upregulation in IL-10 mRNA in response to *S. pneumoniae* infection (data not shown). These findings show that, in contrast to wild type PMNs, CD73-deficient PMNs produce IL-10 in response to *S. pneumoniae*, and that the induction of IL-10 production upon infection can be blocked by EAD.

**Figure 4.**
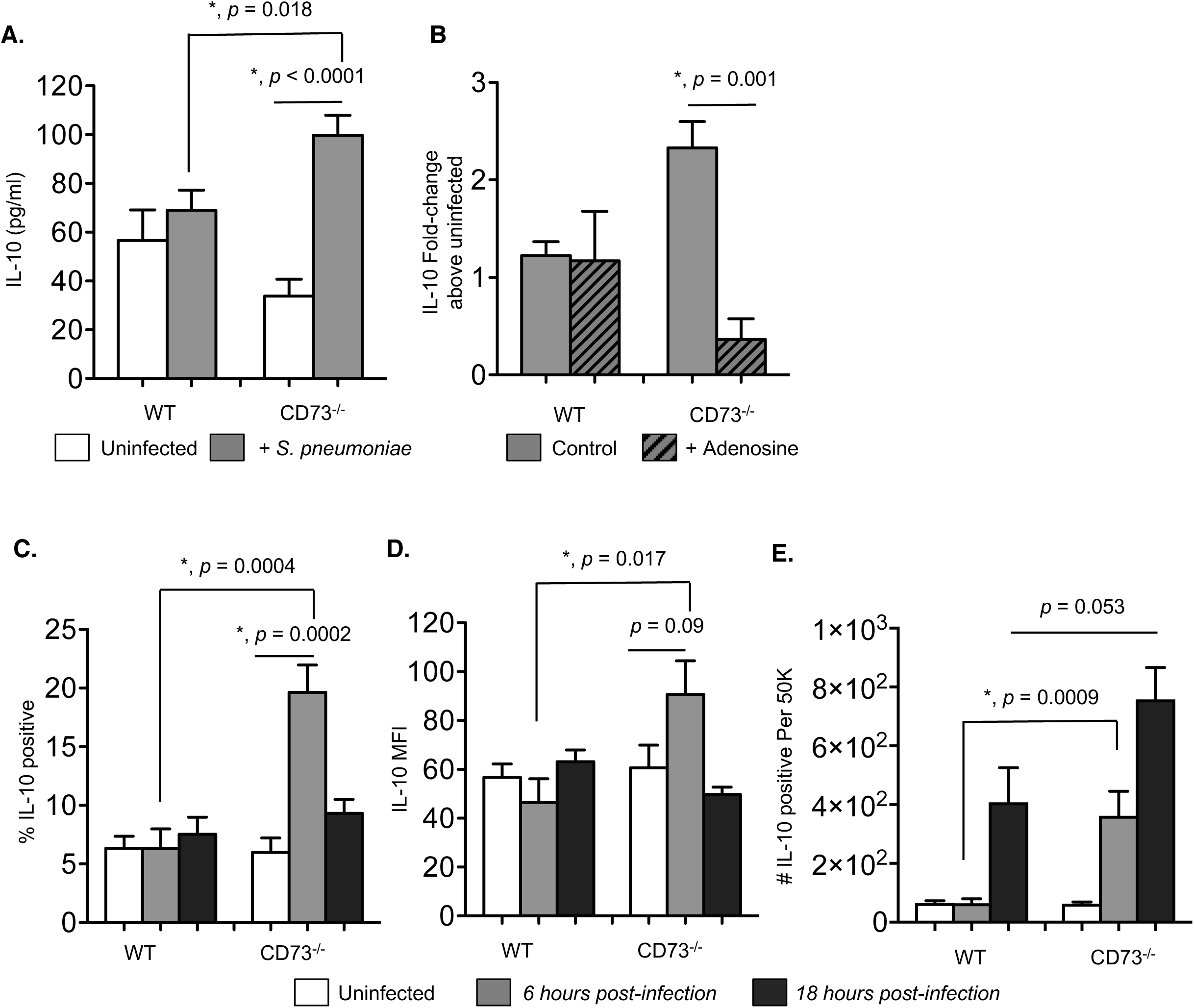
CD73 inhibits IL-10 production from PMNs following pneumococcal infection. (A) PMNs from the indicated mouse strains were incubated for 45 minutes at 37°C with *S. pneumoniae* pre-opsonized with homologous sera or mock treated with buffer and homologous sera (uninfected) *in vitro*. The supernatants were then collected and assayed for IL-10 production by ELISA. (B) PBS control or adenosine (100μM) were added to the PMNs 30 minutes prior to *in vitro* infection and the fold-change in IL-10 production was calculated by dividing the values of infected reactions by uninfected controls for each condition. Data were pooled from three separate experiments (n=3 mice) with each condition tested in triplicate per experiment. Asterisks indicate significant differences determined by Student’s t-test. (C-E) Wild-type C57BL/6 or CD73^-/-^ mice were mock-infected or i.t challenged with 5 x 10^5^ CFU of *S. pneumoniae*. Six (grey bars) and 18 hours (black bars) following challenge, (C) the percentage of IL-10 producing PMNs (Ly6G+), (D) the mean florescent intensities (MFI) of IL-10 in PMNs (Ly6G+) and (E) the number of IL-10 producing PMNs (Ly6G+) recruited into the lungs were determined by intracellular cytokine staining and flow cytometry (see Materials and Methods). Pooled data are from three separate experiments (n=6-9 mice per strain per time point). Significant differences determined by Student’s t-test are indicated by asterisks.

### 3.5 Pulmonary PMNs from CD73^-/-^ mice produce IL-10 early on during *S. pneumoniae* infection

Next, we wanted to test the relevance of our findings *in vivo* and assess IL-10 production during lung infection. WT B6 and CD73^-/-^ mice were infected i.t. with 5×10^5^ CFU of *S. pneumoniae* TIGR4 and IL-10 production by pulmonary PMNs was monitored following infection by intracellular cytokine staining. We focused on the first 18 hours of infection since that is when PMNs are most relevant for the control of bacterial numbers. When we gated on PMNs (Ly6G+; see Fig. S3 for the gating strategy), we found that a low percentage (∼6%) of PMNs in the lungs of WT mice expressed IL-10 and similar to our observations *in vitro*, there was no increase in IL-10 production upon infection (Fig. 4C and D) within the first 6 hours. However, at this timepoint, we observed a significant increase in IL-10 production by pulmonary PMNs in CD73^-/-^ mice. The percentage of IL-10 producing CD73^-/-^ PMNs increased from 5% at baseline to 20% (Fig. 4C) and the amount of IL-10 produced (measured by MFI) increased approximately 1.5-fold (Fig. 4D). When we compared the number of PMNs producing IL-10, we found that in CD73^-/-^ mice, there is a significant increase in IL-10 producing PMNs above baseline by 6 hours post-infection. This increased IL-10 production by CD73^-/-^ PMNs was not due to higher bacterial numbers, as we had previously found that at 6 hours post-infection bacterial burdens in WT and CD73^-/-^ mice were equivalent and differences in bacterial control became apparent after that timepoint (3).

In contrast, in WT mice, we did not observe an increase in IL-10 producing PMNs until 18 hours following infection (Fig 4C-E). Further, as compared to CD73^-/-^ mice, WT controls had 6-fold and 2-fold fewer PMNs making IL-10 at 6 and 18 hours after lung challenge, respectively (Fig 4E). When we measured total IL-10 production in the lungs at 6 hours following infection, we found that CD73^-/-^ mice had on average more IL-10 (231.9 +/-20.51 pg/ml) as compared to WT mice (190 +/-86.65 pg/ml), but the difference did not reach statistical significance. These findings show that upon lung infection, in the absence of CD73, IL-10 production by pulmonary PMNs is higher and occurs more rapidly.

### 3.6 Blocking IL-10 prior to infection rescues the susceptibility of CD73^-/-^ mice to pneumococcal lung infection

To test whether the early increase in IL-10 production by PMNs in CD73^-/-^ mice contribute to their susceptibility to infection, we treated the mice with either isotype control or an IL-10 blocking antibody 2 hours prior to infection. Mice were then challenged i.t. with 5×10^5^ CFU of *S. pneumoniae*, and bacterial burdens in the lung and blood were determined two days post-infection. Blocking IL-10 prior to infection significantly boosted the resistance of CD73^-/-^ mice resulting in a 20 and 24-fold reduction in bacterial numbers in the lung and blood respectively as compared to isotype treated controls (Fig 5A and B). Importantly, blocking IL-10 in CD73^-/-^ mice rendered bacterial burdens in these mice indistinguishable from those of WT mice (Fig 5A and B). In WT mice, blocking IL-10 prior to infection had no significant effect on bacterial burden (Fig 5A and B). Our data suggest that the increased susceptibility of CD73^-/-^ mice during pneumococcal infection is in part mediated by increased IL-10 production.

**Figure 5.**
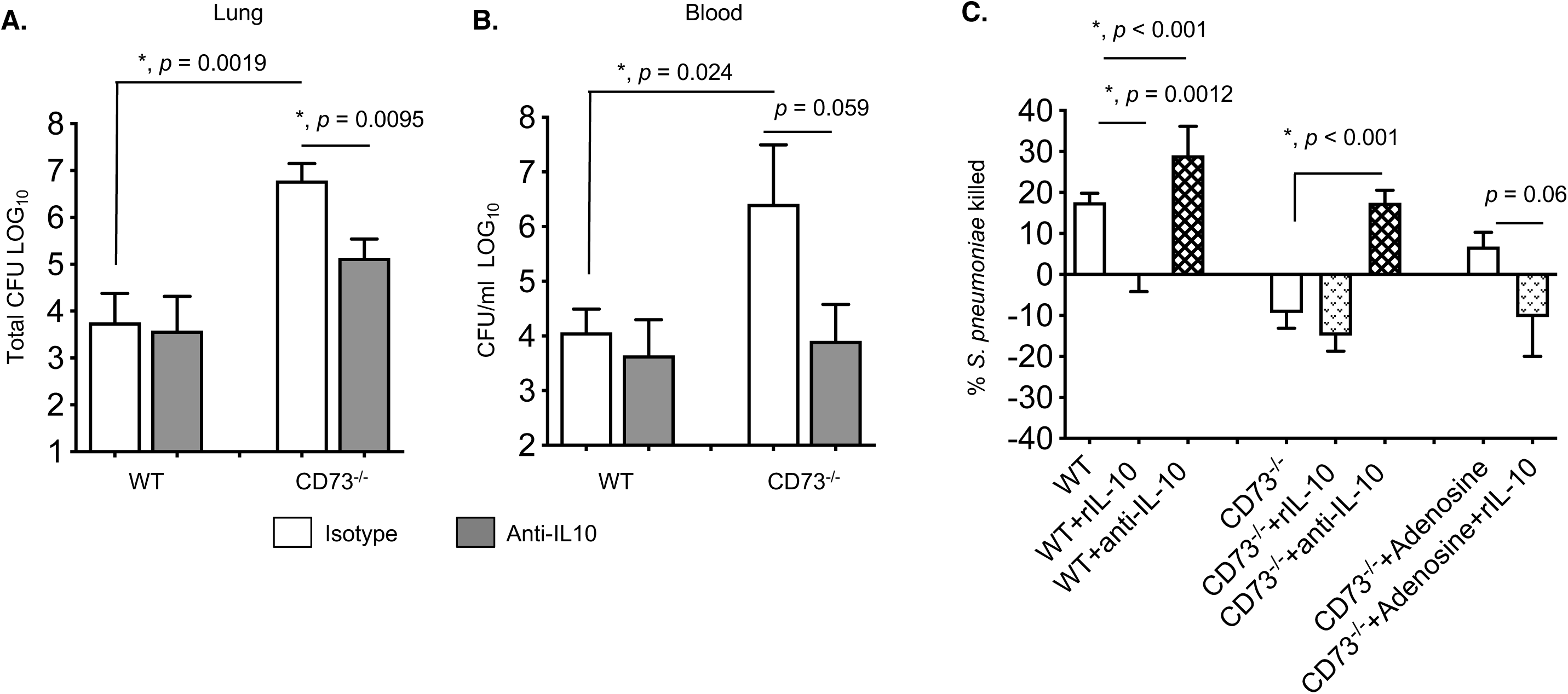
Blocking IL-10 rescues the anti-microbial function of CD73^-/-^ PMNs and reverses the susceptibility of CD73^-/-^ mice to pneumococcal challenge. (A-B) Wild-type (WT) C57BL/6 or CD73^-/-^ mice were treated i.p with IL-10 blocking antibody JES5-2A5 (anti-IL-10) or isotype control two hours prior to pulmonary challenge with 5×10^5^ CFU of *S. pneumoniae*. Pneumococcal burdens in the lungs (A) and blood (B) were determined 48 hours post-infection. Pooled data from three separate experiments with (CD73^-/-^ +/-anti-IL-10 n=8 mice; WT+ anti-IL-10 n=5 mice and WT + isotype n=13 mice per group) are shown. Values significantly different by Student’s t-test are indicated by asterisk. (C) PMNs from C57BL/6 or CD73^-/-^ mice were incubated for 20 minutes with the indicated anti-IL-10 (1μg/ml JES5-2A5), isotype control (1μg/ml), rIL-10 (50ng/ml) or adenosine (100μM) and then infected with pre-opsonized *S. pneumoniae* for 45 minutes at 37°C. Reactions were then plated on blood agar plates and the percentage of bacteria killed compared to a no PMN control under the same conditions was calculated. Data shown are pooled from three separate experiments (n=3 mice per strain) with each condition tested in triplicate. Asterisks indicate significant differences determined by Student’s t-test.

### 3.7 IL-10 impairs the ability of PMNs to kill *S. pneumoniae*

Although the role of IL-10 during *S. pneumoniae* infection in mice is well established (10–12, 38), its effect on PMN anti-pneumococcal function has not been elucidated. To test that, WT PMNs were treated with either recombinant IL-10 or IL-10 blocking antibody and their ability to kill pneumococci measured in our *in vitro* opsonophagocytic assay. We found that addition of IL-10 abrogated the ability of WT PMNs to kill pneumococci while blocking this cytokine slightly boosted the anti-bacterial efficiency of PMNs (Fig 5C).

To test whether the anti-bacterial activity of CD73^-/-^ PMNs can be rescued by targeting IL-10, we treated the PMNs with the IL-10 blocking antibody and performed the opsonophagocytic assay. We found that blocking IL-10 restored the pneumococcal killing ability of CD73^-/-^ PMNs to WT levels (Fig 5C). To test if the ability of adenosine to rescue CD73^-/-^ PMN function was dependent on inhibiting IL-10, we added recombinant IL-10 to CD73^-/-^ PMNs supplemented with adenosine. We found that the addition of recombinant IL-10 had no discernable effect on bacterial killing by CD73^-/-^ PMNs, consistent with the (already high) levels of IL-10 produced by these cells (Fig 5C). However, the addition of IL-10 prevented the ability of adenosine to boost the anti-microbial function of these cells (Fig 5C). These findings demonstrate that EAD’s ability to inhibit IL-10 production is crucial for the anti-pneumococcal function of PMNs.

### 3.8 CD73 has no effect on PMN viability, bacterial uptake, intracellular killing or enzymes in primary granules

We next wanted to identify how CD73, EAD and IL-10 were regulating the ability of PMNs to kill *S. pneumoniae*. Bacterial uptake was previously shown to be important for killing by PMNs (4, 5). To test if EAD production was affecting bacterial phagocytosis, we used a gentamicin-protection assay where gentamicin was added following PMN infection with opsonized pneumococci to kill any extracellular bacteria. We found that the pneumococcal pore forming toxin pneumolysin (PLY) underestimated the true numbers of intracellular bacteria (Fig. S4A) in the gentamicin protection assays, but had no effect on susceptibility to the antibiotic (Fig. S4B) or overall opsonophagocytic killing by PMNs as previously described (39) (Fig. S4C). We therefore proceeded with using *S. pneumoniae* Δ*PLY* to compare phagocytosis between WT and CD73^-/-^ PMNs. To differentiate uptake from intracellular killing, we looked early at 10 minutes post infection, and found no significant difference in the percentage of engulfed bacteria between the different mouse strains (Fig. S4D). We also found that the engulfed bacteria were very efficiently killed by both WT and CD73^-/-^ PMNs (Fig. S4E). We observed 50% and 100% of the engulfed inoculum being killed by PMNs at 30 and 90 minutes post-uptake respectively. We also tested the effect of IL-10 on the association of PMNs with GFP-expressing pneumococci and found that neither addition nor blocking of IL-10 altered association (data not shown). These findings suggest that CD73/EAD do not affect bacterial uptake or intracellular killing by PMNs.

Components of PMN primary granules including serine protease such as cathepsin G (CG) and neutrophil elastase (NE) (4, 5) as well as myeloperoxidase (MPO) (40) were shown to be important for the ability of PMNs to kill *S. pneumoniae*. We therefore measured if CD73, EAD and IL-10 altered the activities of serine proteases Cathepsin G and NE. We were unable to detect Cathepsin G enzymatic activity either at baseline or upon pneumococcal infection in any of the samples we collected. NE activity was easily detected; however, we found no significant difference in NE activity between WT and CD73^-/-^ PMNs at baseline or upon infection. Further, addition of adenosine, recombinant IL-10 or the IL-10 blocking antibody did not alter this response (Fig S5A). When we measured MPO levels, we found that pneumococcal infection significantly increased the amounts of intracellular and released MPO (Fig S5B and C) as previously reported (40). However, there was no difference in the amount of MPO between WT and CD73^-/-^ PMNs, nor were its levels altered by blocking or adding IL-10 (Fig S5B and C). These findings show that CD73, EAD and IL-10 had no effect on the primary granule components NE or MPO.

It was previously reported that IL-10 producing PMNs were apoptotic (41). Since cellular death pathways including apoptosis and necrosis are known to play a role in the anti-bacterial function of PMNs (42), we compared the percentage of apoptotic and necrotic cells using Annexin V and propidium iodide staining and flow cytometry. We found that as expected (43), pneumococcal infection induced pore formation and necrosis of PMNs (PI and Annexin V double positive cells) (Fig S6). However, there was no difference in viability of PMNs at baseline or upon infection between WT and CD73^-/-^ PMNs. Addition or inhibition of IL-10 did not further alter the percentage of necrotic PMNs (Fig S6). These findings suggest that CD73, EAD and IL-10 do not affect PMN viability.

**Figure 6.**
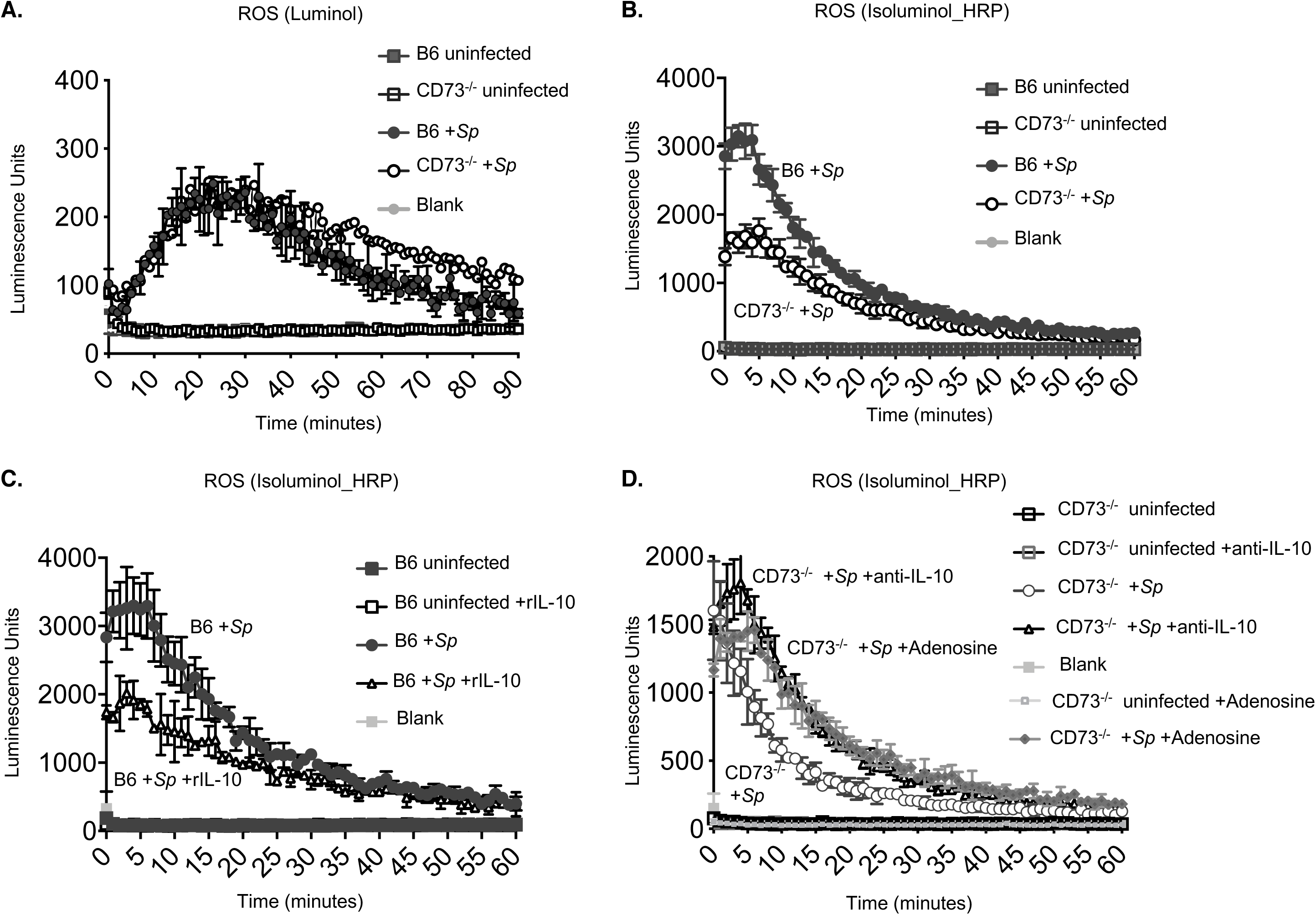
Inhibition of IL-10 production by CD73 is important for optimal production of reactive oxygen species by PMNs in response to *S. pneumoniae* infection. PMN were isolated from the bone marrow of the indicated strains of mice and left untreated (A and B) or treated for 30 minutes with the indicated rIL-10 (50ng/ml) (C) or 100μM Adenosine or anti-IL-10 (1μg/ml JES5-2A5) or isotype control (1μg/ml) (D). PMNs were then infected with *S. pneumoniae* pre-opsonized with homologous sera (+*Sp*) or treated with 3% matching sera (uninfected) and intracellular ROS production measured by chemiluminescence of Luminol (A) or extracellular ROS production measured by chemiluminescence of Isoluminol in the presence of HRP (B-D). Representative data are shown from one of six (B) and one of four (A, C-D) separate experiments with one mouse per strain per experiment where each condition is tested in quadruplicates. Significant differences (*p<0.05*) were determined by 2-way ANOVA followed by Sidak’s multiple comparisons test.

### 3.9 CD73 and IL-10 regulate extracellular ROS production by PMNs upon *S. pneumoniae* infection

IL-10 was reported to down-regulate reactive oxygen species (ROS) production by PMNs (18). To determine whether the CD73/IL-10 axis regulated ROS production upon pneumococcal infection, we first compared production of ROS by WT and CD73^-/-^ PMNs. We first measured ROS levels using luminol, which is cell permeable and detects intracellular ROS. We found that the infection up regulated ROS production rapidly within minutes to more than 50-fold increase around the peak of the response at the 25 minute mark in both strains of mice (Fig. 6A). There was no significant difference in intracellular ROS levels by WT vs CD73^-/-^ PMNs in response to pneumococcal infection, consistent with our findings that bacterial uptake and intracellular killing was not affected by CD73.

As PLY forms pores that can lead to the release of products from PMNs, we next examined levels of ROS using isoluminol and HRP which can detect extracellular ROS. In uninfected PMNs, no ROS production was observed (Fig. 6B). However, upon exposure to pneumococci, ROS was rapidly induced by PMNs from both strains of mice. This response was immediate and peaked within the first five minutes of infection and waned over time (Fig. 6B) as previously described (24). Importantly, during the first 10 minutes, PMNs from B6 mice released significantly higher levels of extracellular ROS than CD73^-/-^ PMNs, producing 3-fold higher levels of ROS at the peak of the response (Fig. Fig. 6B at the 5 minute mark). Extracellular ROS levels by PMNs in response to the positive control PMA was not different between mouse strains (not shown), indicating that the blunted response observed in CD73^-/-^ PMNs was not due to a general defect in ROS production, but rather was specific to pneumococcal infection.

To determine if IL-10 had a role in production of ROS by PMNs, we added recombinant IL-10 to B6 PMNs. We found that this cytokine significantly blunted the magnitude of the ROS response during pneumococcal infection, which is consistent with its reported anti-inflammatory role (18) (Fig. 6C). Next we tested whether ROS production by CD73^-/-^ PMNs upon infection can be rescued by treating the PMNs with the IL-10 blocking antibody. We found that blocking IL-10 significantly boosted extracellular ROS levels in infected CD73^-/-^ PMNs, resulting in around a 2-fold increase at the peak of the response at 5 minutes post infection (Fig. 6D). Similarly, addition of adenosine significantly boosted ROS responses by infected CD73^-/-^ PMNs (Fig. 6D). Taken together, these data demonstrate that inhibition of IL-10 production by CD73 is important for an optimal ROS response by PMNs during *S. pneumoniae* infection.

### 3.10 ROS detoxification is important for the ability of *S. pneumoniae* to resist killing by WT but not CD73^-/-^ PMNs

The role of ROS production by NADPH oxidase in the function of PMNs against pneumococci is unclear (32, 44). Consistent with previous reports (4), when we treated PMNs with DPI, which inhibits ROS production by flavoenzymes such as NADPH oxidase, we found no effect on the ability of these cells to kill *S. pneumoniae* (Fig 7A). However, when we used ascorbic acid, an anti-oxidant that scavenges ROS (45) or EUK 134 which detoxifies ROS by mimicking the activity of superoxide dismutase and catalase (46), the ability of WT PMNs to kill *S. pneumoniae* was completely abrogated (Fig 7A). We also tested *S. pneumoniae* that lack manganese-dependent superoxide dismutase (Δ*sodA*), the major SOD enzyme in *S. pneumoniae* that degrades superoxide radicals (47). We found that this bacterial mutant was significantly more susceptible to killing by PMNs from C57BL/6 mice as compared to wildtype *S. pneumoniae* (Fig 7A and B). Addition of EUK 134 or ascorbic acid reversed the susceptibility of Δ*sodA S. pneumoniae* to killing by PMNs (Fig 7A). None of the compounds tested had a direct effect on bacterial viability (not shown). These findings suggest that *S. pneumoniae* are susceptible to killing by ROS produced by PMNs.

**Figure 7.**
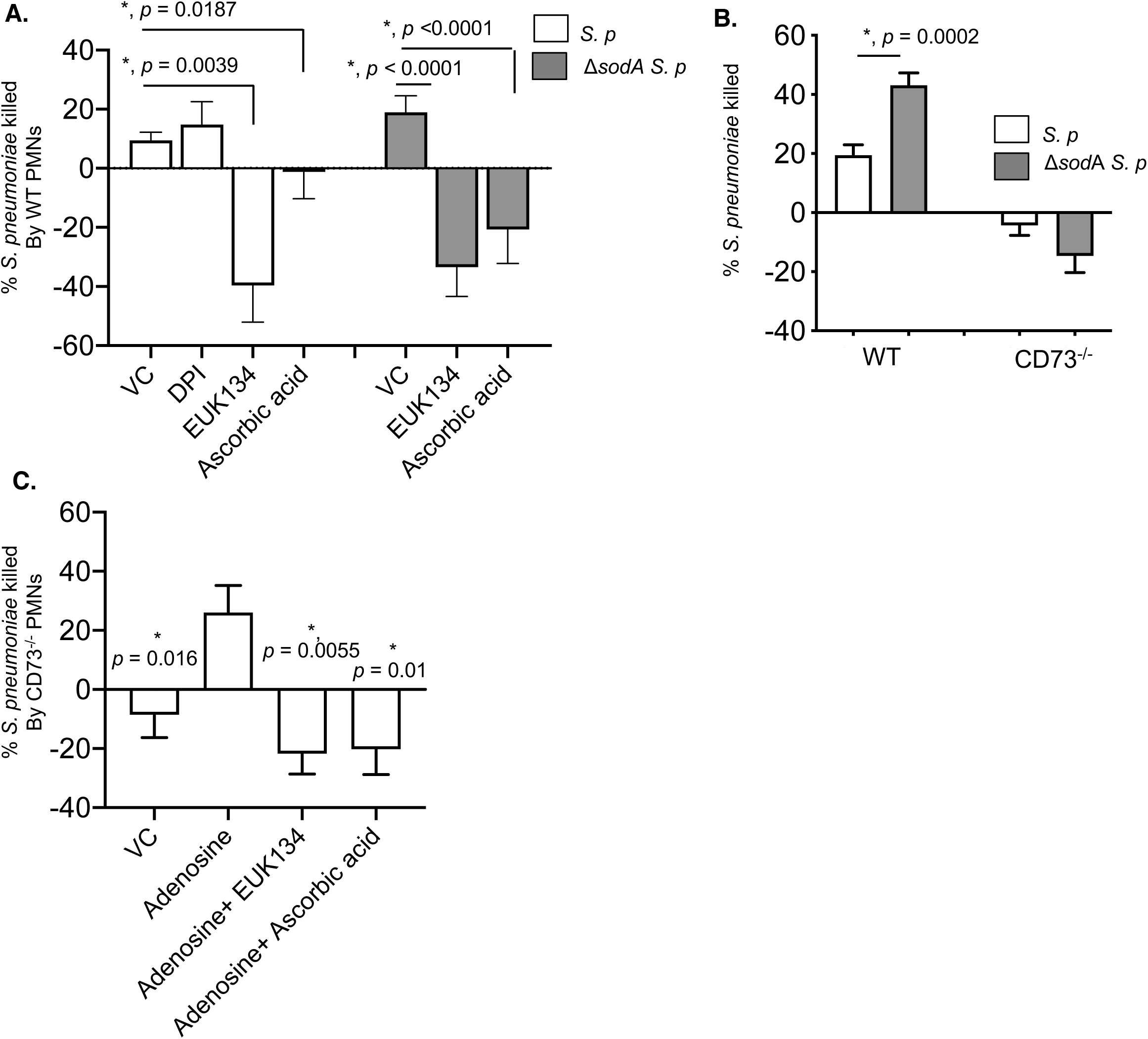
Detoxification of ROS is important for *S. pneumoniae* to resist killing by wildtype, but not CD73^-/-^ PMNs. (A) PMNs isolated from the bone marrow of C57BL/6 (WT) mice were treated with PBS (VC), 10μM DPI, EUK 134 (25μM) or Ascorbic acid (100μM) for 30 minutes at 37°C. (B) PMNs isolated from the bone marrow of C57BL/6 (WT) or CD73^-/-^ mice were left unmanipulated. (C) PMNs isolated from the bone marrow of CD73^-/-^ mice were treated with 100μM adenosine in the absence or presence of EUK 134 or Ascorbic acid for 30 minutes at 37°C. (A-C) The PMNs were then infected with the indicated wild type or *ΔsodA S. pneumoniae* in the presence of homologous sera for 45 minutes at 37°C. Reactions were then stopped by placing samples on ice and viable CFU were determined after serial dilution and plating. The percentage of bacteria killed by comparing surviving CFU to a no PMN control under the same conditions. Data shown are pooled from four separate experiments (n=4 biological replicates or mice per strain) where each condition was tested in triplicate (n=3 technical replicates) per experiment. Asterisks indicate significance calculated by Student’s t-test.

When we tested CD73^-/-^ PMNs, we did not observe any increase in killing of Δ*sodA S. pneumoniae* (Fig 7B), which was expected given the blunted ROS response by these PMNs. To test if the presence of ROS is important for the ability of adenosine to rescue CD73^-/-^ PMN function, we used EUK 134 or ascorbic acid. We found that the ability of adenosine to boost the anti-pneumococcal activity of these cells was lost when ROS was removed (Fig 7C). These findings suggest that the presence of ROS is important for EAD’s ability to promote the anti-pneumococcal function of PMNs.

## 4. Discussion

There is mounting evidence that PMNs are plastic and display phenotypic and functional heterogeneity under different disease states (48). These range from traditional pro-inflammatory subsets during infections (13, 48) to B-helper PMNs (32, 49) to suppressive phenotypes in tumor microenvironments (50, 51) and anti-inflammatory in response certain pathogens and microbial products (13-16, 52-54). In the context of pneumococcal pneumonia, we previously found that the efficacy of PMNs during the course of disease changes from clearing bacteria early on to promoting infection at later time points (3). Here we show that this shift away from an anti-microbial phenotype is associated with a decrease in CD73 expression on PMNs. Using *in vitro* bacterial killing assays and adoptive transfer of PMNs into CD73^-/-^ recipients, we found that CD73 expression on PMNs was crucial for the ability of these cells to kill *S. pneumoniae* and mediate protection during lung infection *in vivo.* In fact, loss of CD73 expression on PMNs seem to actively promote bacterial growth as *S. pneumoniae* grew in the presence of CD73^-/-^ PMNs *in vitro* and adoptive transfer of CD73^-/-^ PMNs worsened the infection resulting in increased bacterial burdens in recipient mice. Importantly, inhibition of CD73 *in vitro* abrogated the ability of human PMNs to kill pneumococci in 60% of donors tested, highlighting the clinical relevance of this pathway. These findings suggest that CD73 regulates the PMN antimicrobial phenotype.

During systemic infection with *S. pneumoniae,* a subset of immature PMNs with a lower expression of surface Ly6G that exhibited a low ability to bind bacteria was observed immobilized in the spleen (32). When we followed Ly6G expression on PMNs (gated on Ly6G positive cells) over time, we noticed it changed over time where we observed an increase in Ly6G expression on pulmonary PMNs recruited at the 6 hour time point followed by a subsequent decrease after that. However, we observed that CD73-negative PMNs in the lungs had significantly lower amounts of Ly6G on their surface as compared to their CD73^+^ counterparts at all time points suggesting that they could denote a less mature subset. We believe that these CD73-negative pulmonary PMNs migrate from the circulation after originating from the bone marrow, as the reduced expression of CD73 following infection was also observed on circulating PMNs as well as the PMN pool in the bone marrow.

In exploring mechanisms of how CD73 controls the PMN antimicrobial phenotype, we found that it inhibited IL-10 production in response to pneumococcal infection. In wild type mice, we did not see any up regulation in IL-10 production by PMNs early in infection. IL-10 producing PMNs only started to accumulate at the 18h time point that coincided with a decrease in CD73 on pulmonary PMNs and their inability to control bacterial numbers (3). In contrast, PMNs from CD73^-/-^ mice up regulated IL-10 production early on within 6 hours of infection. Our IL-10 blocking experiments both *in vitro* and *in vivo* suggest that this early IL-10 production impaired the ability of CD73^-/-^ mice to control infection. Our data align with previous reports that administration of IL-10 early in infection impairs the ability of the host to control pneumococcal numbers (10). However, studies have also found that mice that lack IL-10 had exacerbated PMN influx in their lungs (12, 38) and succumbed to the infection despite having lower bacterial burden in their organs (12). These findings suggest that early production of IL-10 is deleterious for the host’s ability to control the infection, however, it may be required later on for resolution of inflammation and return to homeostasis. Therefore, the eventual accumulation of IL-10 producing PMNs in wild type mice we observed here may be beneficial in resolution of pulmonary inflammation observed in healthy hosts (3).

Murine PMNs have been described to produce IL-10 in sepsis models and in response to several bacterial infections such as *Escherichia coli*, *Shigella flexneri* (16) and *Staphylococcus aureus* (13) as well as parasitic infections such as *Leishmania major* (14) and *Trypanosoma cruzi* (15). In contrast, IL-10 production by pulmonary PMNs was actively repressed during *Mycobacterium tuberculosis* (55). In this study, we did not detect an up regulation in IL-10 production by PMNs in response to *S. pneumoniae* infection in wild type mice either *in vivo* during lung challenge or *in vitro* in response to direct bacterial infection. Further, we were unable to detect an increase in IL-10 mRNA in PMNs isolated from human donors in response to stimulation with *S. pneumoniae*. This fits with previous studies demonstrating that resting human PMNs have histone modifications at the IL-10 locus rendering it transcriptionally silent (56) and that induction of IL-10 production by PMNs requires further stimulation such as direct contact with LPS-treated Tregs or IL-10 itself that promote histone posttranslational modifications that can activate IL-10 transcription (41).

Our data suggests that IL-10 production by PMNs in response to *S. pneumoniae* is actively suppressed by EAD production by CD73. This is in contrast to what has been described for macrophages. Adenosine induced IL-10 production by macrophages in response to LPS stimulation (36, 37) or *E. coli* infection (35) by acting via the A2A or A2B receptor to facilitate binding of transcription factors to the IL-10 locus and enhancing transcription (35) or via posttranscriptional modifications of the 3’-UTR that facilitated translation (36) respectively. These differences observed could be accounted for by the cell type i.e. macrophages vs. PMNs or the stimulation where the macrophage studies have been performed with Gram-negative bacteria and their products (LPS) while we are examining responses to a Gram-positive organism.

Phagocytosis and serine proteases were shown to contribute to the ability of PMNs kill *S. pneumoniae* (4). In exploring how the CD73/IL-10 axis regulated PMN anti-pneumococcal activity we found that apoptosis, bacterial uptake and intracellular killing and production of anti-microbial Neutrophil Elastase and Myeloperoxidase were not affected by this pathway. Rather, inhibition of IL-10 production by CD73 was crucial for optimal production of extracellular ROS by PMNs upon *S. pneumoniae* infection. Importantly, the presence of ROS was required for the ability of adenosine to rescue bacterial killing by CD73^-/-^ PMNs. This is in line with previous findings that adenosine signaling via the A1 receptor enhanced superoxide production by PMNs in response to antibody coated erythrocytes (57) and that IL-10 blunted PMN superoxide production in response to PMA and *C. albicans* hyphae (18, 19).

We found here that the bacterial infection triggered ROS production by PMNs similar to what has been reported (24, 58). ROS detoxification is crucial for pneumococcal survival in host environments (59) as bacterial mutants lacking components that detoxify ROS die more readily in response to oxidative stress (60–63). Similarly, we found that Δ*sodA S. pneumoniae*, which lacks the enzyme superoxide dismutase, a manganese-dependent enzyme that detoxifies superoxide radicals (47), is more susceptible to killing by PMNs. Importantly, this susceptibility was reversed upon the exogenous addition of compounds that remove ROS. This highlighted the importance of ROS detoxification for the ability of *S. pneumoniae* to resist killing by PMNs. NADPH oxidase is a major source of ROS in PMNs (64). However, when we used DPI to inhibit NADPH oxidase, we saw no effect on bacterial killing by wildtype PMNs. This is in line with previous work showing that mice lacking components of the NADPH oxidase are not more susceptible to pneumococcal lung infection (32, 44) and that using DPI to inhibit respiratory burst has no effect on the ability of PMNs to kill *S. pneumoniae* (4). Intriguingly, when we used EUK134 and ascorbic acid to detoxify or scavenge ROS respectively (45, 46), the ability of PMNs to kill *S. pneumoniae* was significantly impaired. These findings suggest that ROS is important for bacterial killing, however, NADPH oxidase may not be the only source of ROS in PMNs. Cellular ROS can be produced from several sources other than NADPH oxidase including the mitochondrial respiratory chain, lipooxygenases, cyclooxygnases among others (65). In fact, previous work found that the NADPH oxidase inhibitor DPI reduced, but did not completely abrogate ROS production by PMNs in response to pneumococcal infection (40). Pneumococcal infection is known to activate cellular lipooxygenases and cyclooxygenases and these enzymes which are expressed by leukocytes but are not inhibited by DPI (6, 66–69), may act as an alternate sources of ROS.

Adenosine is a well-known regulator of PMN function and is known to regulate PMN recruitment, release of inflammatory cytokines, phagocytic capacity and oxidative burst (34). However, the majority of those studies have been conducted in the context of sterile inflammation or in response to *in vitro* stimulation by inflammatory mediators such as fMLP, LPS, TNF or inert particles (34). The role of the EAD pathway in PMN response to infections is now better appreciated and was shown to play a role in host resistance during pulmonary infections with influenza A virus (70, 71), *Klebsiella pneumoniae* (72) and *S. pneumoniae* (3). Here, we identified the mechanisms by which EAD production by CD73 regulated PMN anti-pneumococcal function and further found it was relevant for the function of PMNs from human donors. This may be relevant for incorporating clinically available adenosine-based drugs to combat pneumococcal pneumonia and other serious lung infections in the future.

## Abbreviations

CFU: Colony Forming Units
CT: Cycle Thresh-hold
EAD: Extracellular Adenosine
I.P: Intra Peritoneal
I.T: Intra Tracheal
MFI: (Mean Fluorescent Intensity)
MOI: Multiplicity of Infection
MPO: Myeloperoxidase
NE: Neutrophil Elastase
OPH: Opsonophagocytic
PLY: Pneumolysin
PMA: Phorbol 12-myristate 13-acetate
PMNs: Polymorphonuclear Leukocytes
Pneumococcus: *Streptococcus pneumoniae*
ROS: Reactive Oxygen Species
SodA: Superoxide Dismutase
WT: Wild Type

## Authorship

NS and JNL conducted research, analyzed data and wrote the paper. MB, EYIT, JHY and SER conducted research, JML provided essential expertise and reviewed the paper. ENBG designed research, conducted research, analyzed data, wrote the paper and had primary responsibility for final content.

## Acknowledgments

We would like to acknowledge Andrew Camilli for bacterial strains and Joan Mecsas for ROS detection protocols. This work was supported by National Institute of Health grant R00AG051784 to ENBG.

---

**Figure S1.**
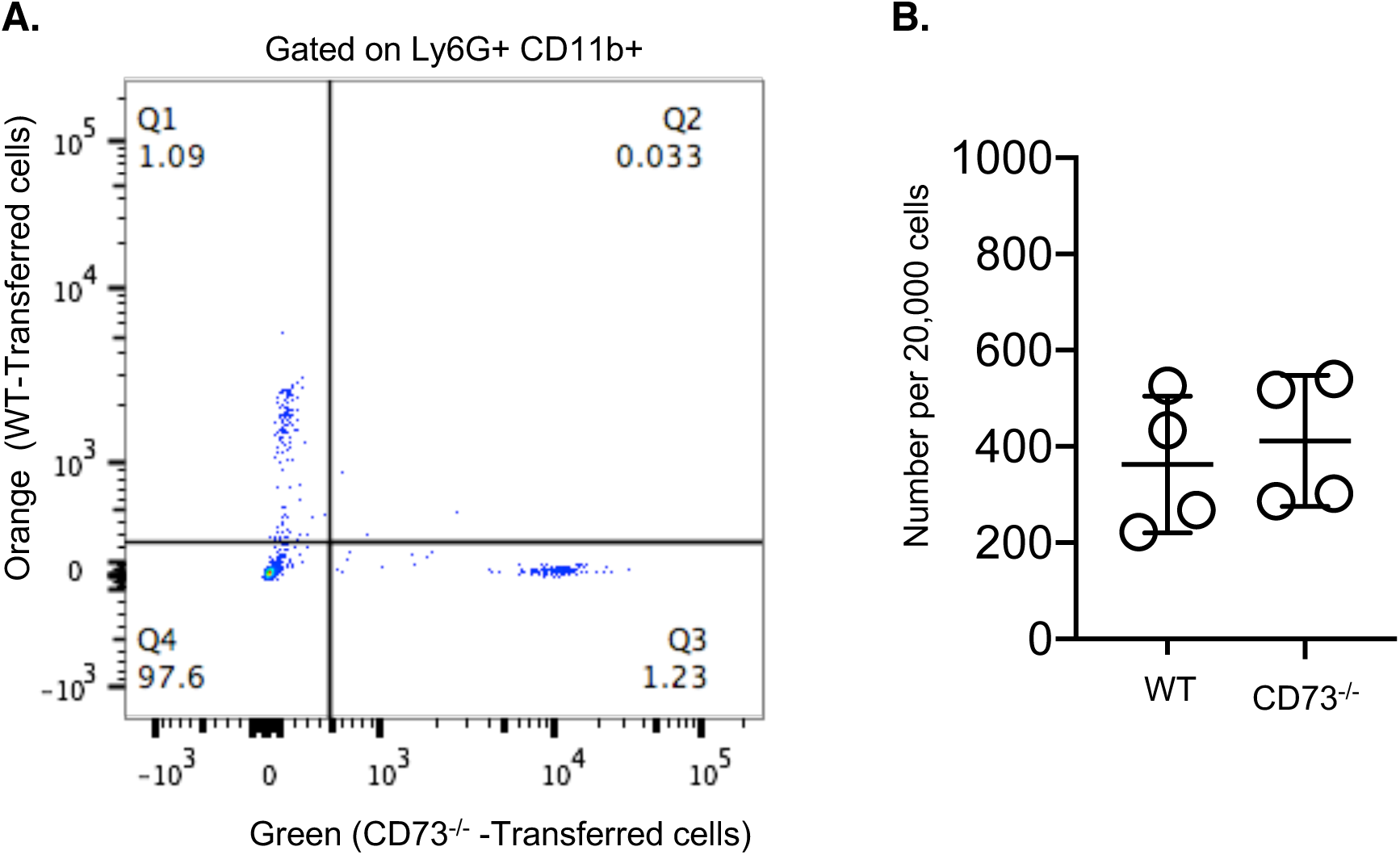
Detection of transferred PMNs in the circulation. CD73^-/-^ mice were adoptively transferred 1×10^6^ PMNs isolated from bone marrows of wildtype (WT) mice and 1×10^6^ PMNs isolated from CD73^-/-^ controls. The transferred PMNs were pre-labeled with CellTracker dyes Orange and Green for WT and CD73^-/-^ respectively and their presence in blood confirmed by flow cytometry 3 hours post transfer. (A) A representative dot plot from one of four recipient mice and (B) the number of the transferred cells per recipient in the blood are shown.

**Figure S2.**
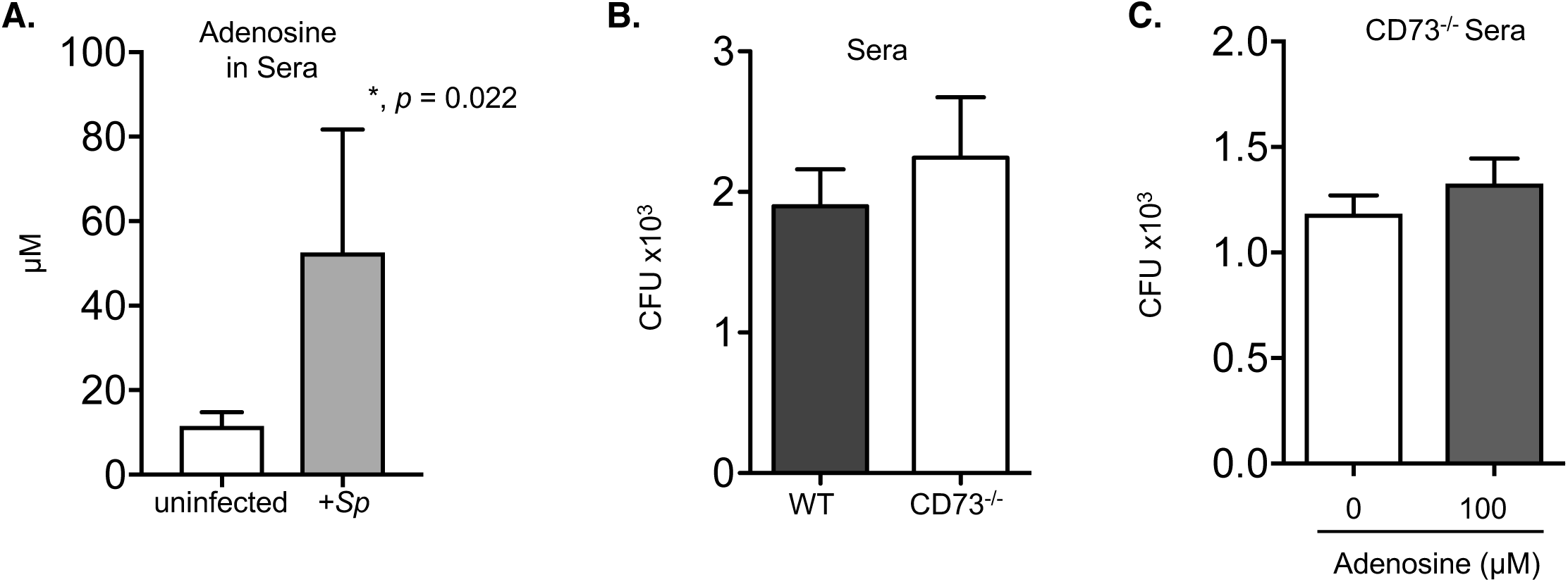
Extracellular adenosine has no direct effect on bacterial survival. (A) The adenosine levels in the sera of mice following pulmonary challenge with 5×10^5^ CFU of *S. pneumoniae* were measured using a fluorometric adenosine assay kit. Data shown are pooled from n=4 uninfected mice and n=7 infected mice. (B) *S. pneumoniae* were incubated with sera only from C57BL/6 (WT) or CD73^-/-^ mice for 40 minutes at 37°C. (C) The indicated concentrations of adenosine or PBS (0) were added prior to the start of incubation. Viable bacteria were enumerated by plating on blood agar plates. (B-C) Data shown represent the means +/-SD and are pooled from two separate experiments with n=2 mice and where each condition was tested in triplicates per experiment.

**Figure S3.**
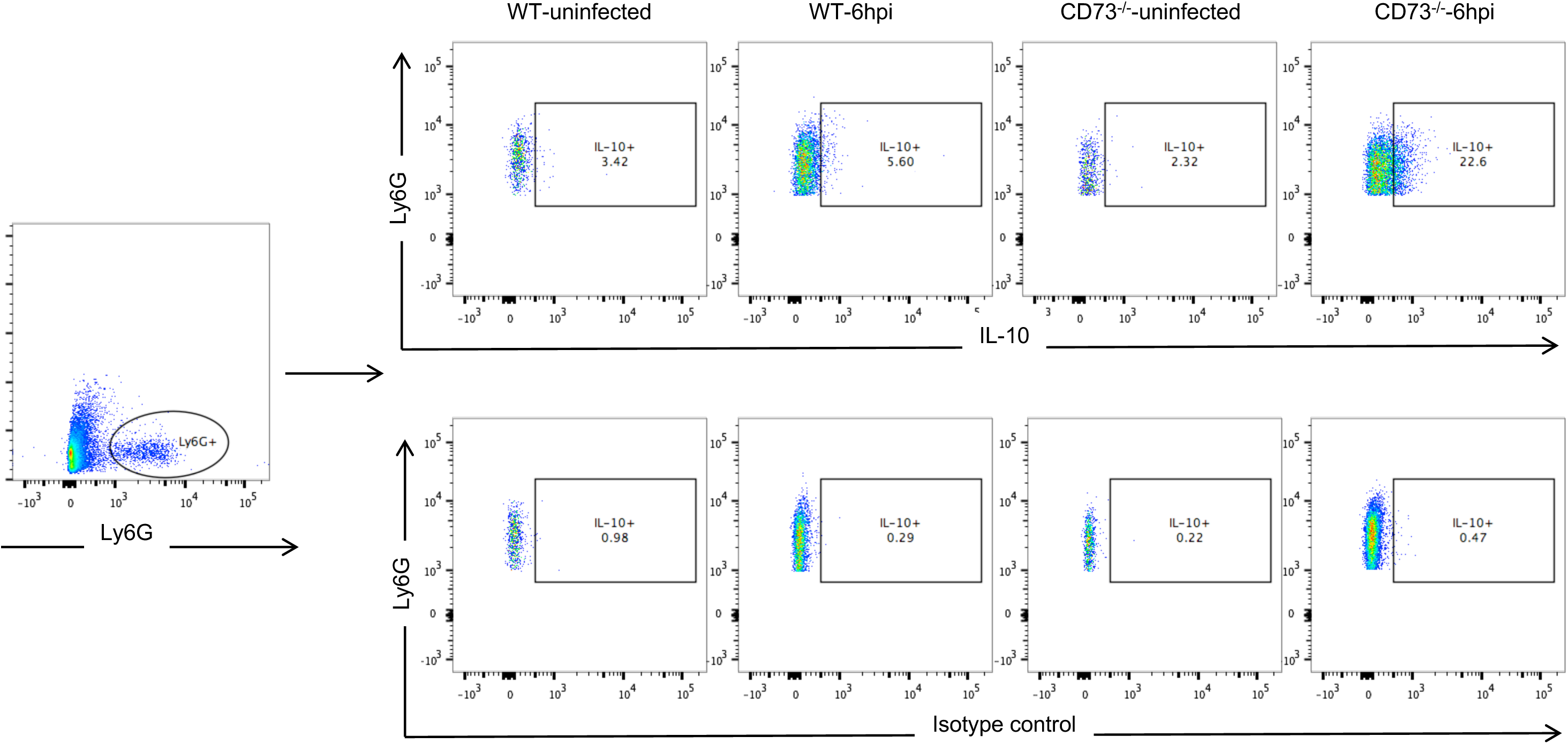
Gating Strategy of IL-10 producing PMNs *in vivo.* Wild-type C57BL/6 or CD73^-/-^ mice were mock-infected or i.t challenged with 5 x 10^5^ CFU of *S. pneumoniae*. Six hours following challenge, the lungs were harvested, incubated with GolgiPlug and stained with Ly6G, IL-10 antibody or isotype control and IL-10 production by PMNs (Ly6G+) and analyzed by flow cytometry. Representative dot plots from one of three separate experiments are shown.

**Figure S4.**
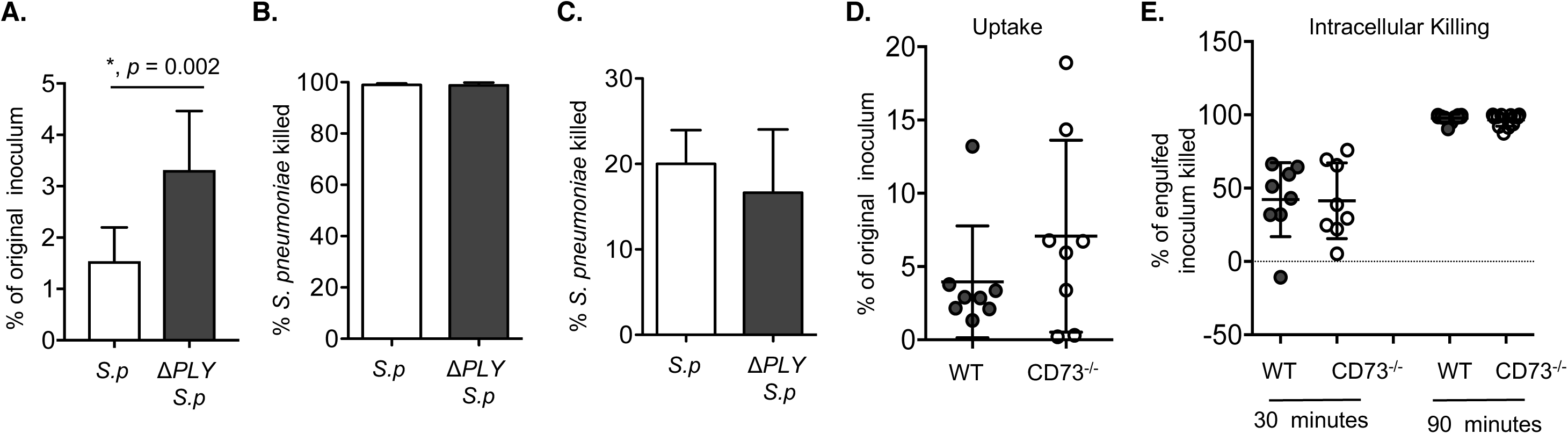
CD73 does not affect bacterial uptake or intracellular killing by PMNs. (A) Bone marrow PMNs from C57BL/6 mice were incubated for 40 minutes at 37°C with wild type or ΔPLY *S. pneumoniae* that were pre-opsonized with homologous sera at an MOI of 2. Gentamicin (50μg/ml) was then added for 30 minutes to kill extracellular bacteria. PMNs were washed and plated on blood agar plates to determine the amount of bacteria engulfed and the % of bacteria taken up (uptake) of the original infecting inoculum was calculated. (B) Wild type *(S.p)* or pneumolysin deficient (*ΔPLY S.p*) *S. pneumoniae* were incubated with 100 μg/ml Gentamycin for 30 minutes and the percentage of bacteria killed compared to a mock treatment control was then calculated by plating on blood agar plates. (C) C57BL/6 PMNs were incubated for 45 minutes at 37°C with the indicated strains of wild type *(S.p)* or pneumolysin deficient (*ΔPLY S.p*) *S. pneumoniae* pre-opsonized with sera. The percentage of bacteria killed compared to a no PMN control under the same conditions was then calculated by plating for viable bacteria on blood agar plates. (D-E) Bone marrow PMNs from the indicated strains of mice were incubated with *S. pneumoniae ΔPLY* pre-opsonized with homologous sera at an MOI of 2 for 10 minutes at 37°C and gentamicin (50μg/ml) was then added for 30 minutes to kill extracellular bacteria. PMNs were then washed and one set (D) immediately plated on blood agar plates to determine the amount of bacteria engulfed and the % of bacteria taken up (uptake) of the original infecting inoculum was calculated. The other sets of PMNs (E) were incubated for 30 and 90 more minutes and then plated to enumerate viable bacteria. The % of bacteria of the engulfed inoculum that was killed was then calculated (Intracellular Killing). Data shown are pooled from (A) two separate experiments, (B) two separate experiments and (C) four separate experiments (n=4 mice) and (D-E) two separate experiments (n=2 mice per strain) where each condition was tested in quadruplicates per experiment. Values significantly different by Student’s t-test are indicated by asterisk.

**Figure S5.**
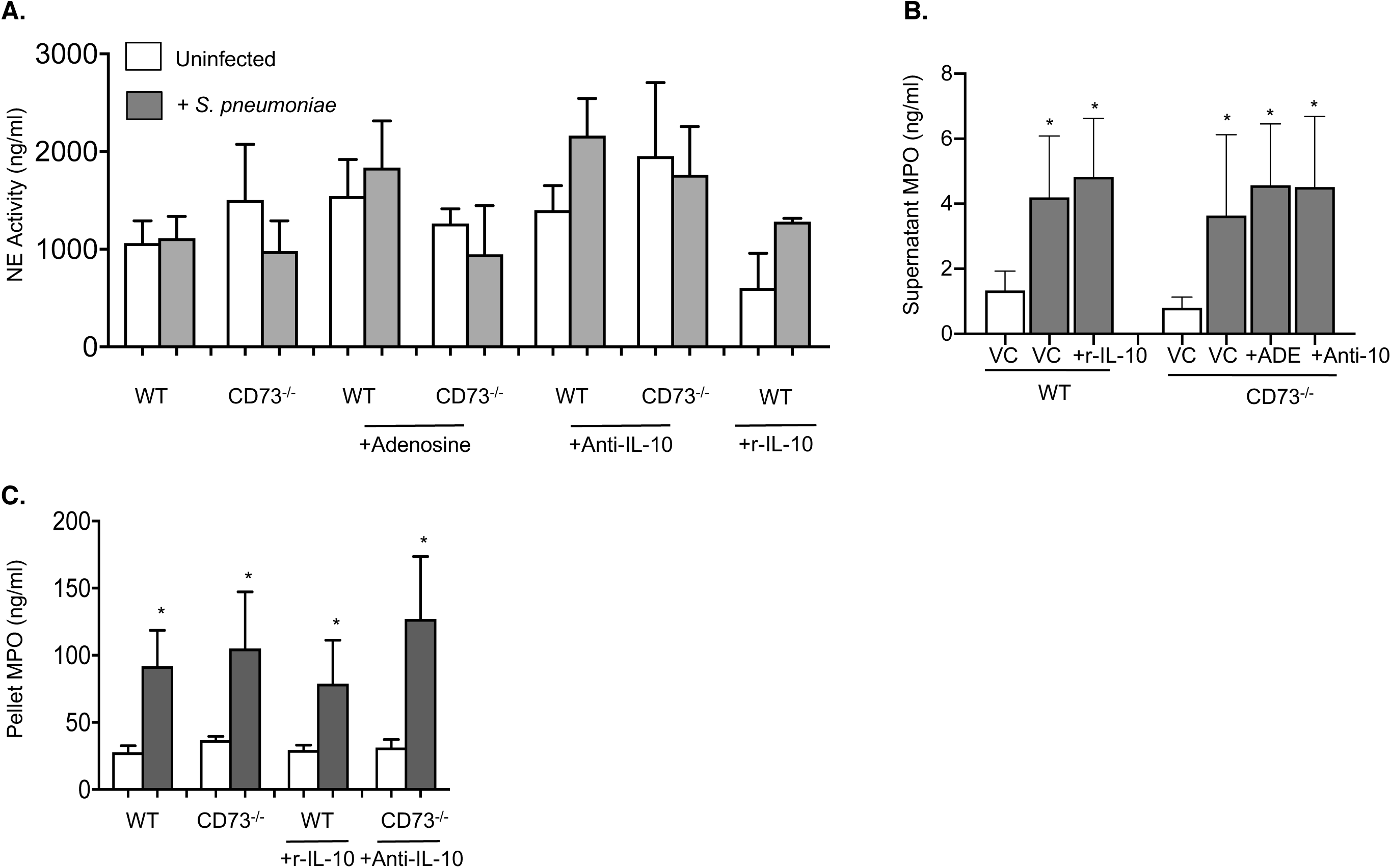
CD73 does not affect production of Neutrophil Elastase or myeloperoxidase. (A) PMNs were incubated for 45 minutes at 37°C with pre-opsonized *S. pneumoniae* or mock treated (uninfected) in the presence of control or adenosine (100μM), anti-IL-10 (1μg/ml JES5-2A5) or rIL-10 (50ng/mL). The supernatants were then collected and assayed for (A) Neutrophil Elastase activity (NE) and (B) myeloperoxidase (MPO) levels. (C) The cells were lysed and assayed for MPO levels. Data were pooled from four separate experiments with 4 mice per strain per condition. Asterisk indicated values significantly different from uninfected controls within the same condition by Student’s t-test (*p<0.05*).

**Figure S6.**
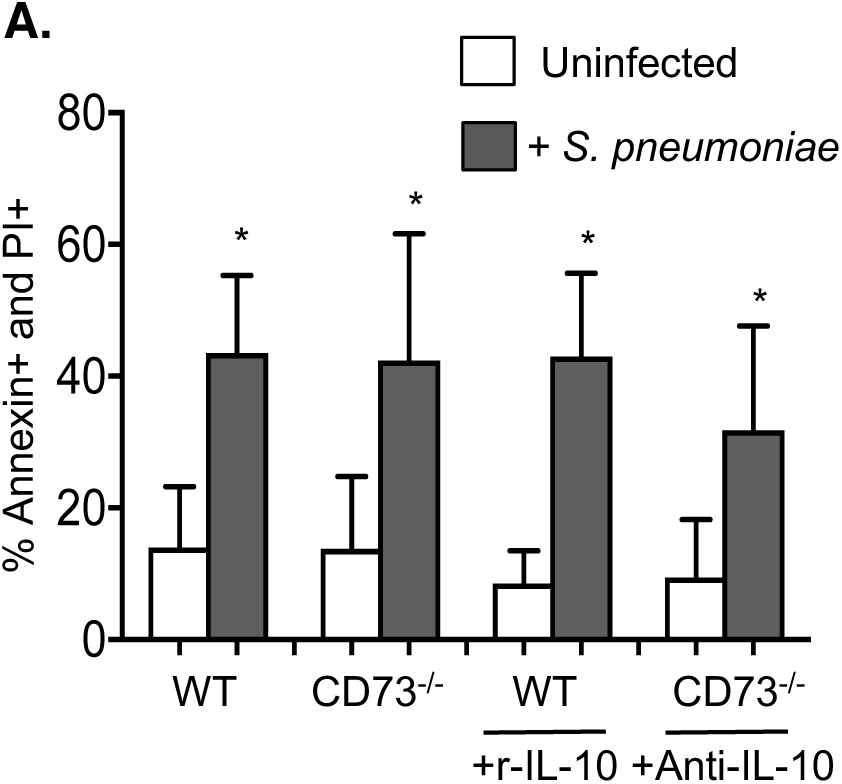
CD73/IL-10 do not impact viability. (A) PMNs were mock treated with PBS or treated for 20 minutes with the indicated anti-IL-10 (1 μg/ml JES5-2A5) or rIL-10 (50ng/ml). PMNs were then incubated for 15 minutes at 37°C with *S. pneumoniae* pre-opsonized with matching sera at an MOI of 2 or mock treated (uninfected) with 3% matching mouse sera only. The percentage of apoptotic cells were then determined by flow cytometry. The percentage of PMNs (gated on Ly6G+ cells) that were double positive for PI and Annexin V are shown. Data were pooled from four separate experiments with 4 mice per strain per condition. Asterisk indicated values significantly different from uninfected controls within the same condition by Student’s t-test (*p<0.05*).

